# Genetic or transcranial magnetic stimulation of B-RAF–MEK signaling promotes CST axon sprouting and functional regeneration

**DOI:** 10.1101/2022.06.01.494346

**Authors:** Francesco Boato, Xiaofei Guan, Yanjie Zhu, Youngjae Ryu, Mariel Voutounou, Christopher Rynne, Chase R. Freschlin, Paul Zumbo, Doron Betel, Katie Matho, Sergey N. Makarov, Zhuhao Wu, Young-Jin Son, Aapo Nummenmaa, Josh Z. Huang, Dylan J. Edwards, Jian Zhong

## Abstract

Facilitating axon regeneration in the injured central nervous system remains a challenging task. RAF–MEK signaling plays an important role in axon elongation during nervous system development. Here we show that activation of B-RAF in mature corticospinal neurons elicited the expression of a discrete set of transcription factors previously implicated in the regeneration of zebrafish optic nerve axons. Genetic activation of B-RAF–MEK signaling promoted robust regeneration and sprouting of corticospinal tract axons after injury. Newly sprouting axon collaterals formed synaptic connections with spinal interneurons, correlating with the recovery of skilled motor function. Seeking a non-invasive way to stimulate axon regeneration, we found that suprathreshold high-frequency repetitive transcranial magnetic stimulation activates the B-RAF canonical effectors MEK1/2 and requires MEK1/2 activity to promote corticospinal axon regeneration and sprouting after injury. These data demonstrate a central role of neuron-intrinsic RAF–MEK signaling in enhancing the growth capacity of mature corticospinal neurons and propose HF-rTMS as a potential therapy for spinal cord injury.

**One Sentence Summary:** Genetic or HF-rTMS-mediated activation of B-RAF– MEK signaling promotes CST axon sprouting and functional regeneration after a spinal cord injury.

## INTRODUCTION

The failure of mature mammalian central nervous system (CNS) neurons to activate cell-intrinsic growth mechanisms and regenerate severed axons contrasts dramatically with the regenerative abilities of mature CNS neurons in lower vertebrates (*1, 2*) or mammalian peripheral neurons (*3*). Identifying ways to engage regenerative axon growth in mature mammalian CNS neurons is a major focus of current research into traumatic CNS injury (*4, 5*). The RAF – MAP kinase signaling cascade mediates long distance axon outgrowth in developing PNS and CNS neurons (*6–10*). This led to the hypothesis that RAF signaling regulates an intrinsic axon growth program and that its activation could drive re-growth of injured corticospinal tract (CST) axons, which are particularly resistant to regeneration (*11–14*). Here, we report that conditional activation of B-RAF signaling in mature corticospinal neurons (CSNs) induces a discrete set of transcription factors (TFs) that overlap substantially with those reported to initiate axon regeneration in zebrafish retinal ganglion cells (RGCs) (Dhara et al., 2019). Several pioneer TFs were found to be regulated by B-RAF signaling. Conditional B-RAF activation in CSNs resulted in significant CST axon sprouting and regeneration in two experimental models of SCI: unilateral pyramidotomy (uPx) and dorsal hemisection (dHx). We present evidence of improved motor functional recovery following uPx and dHx, as well as *de novo* synaptogenesis of newly sprouting CST collaterals. Finally, we explored transcranial magnetic stimulation (TMS) as a clinically applicable approach to non-invasively activate the RAF – MEK signaling in CSNs. Non-invasive brain stimulation has been emerging as a promising strategy to improve function and promote recovery in spinal cord injury (*15–17*), but the underlying plasticity mechanisms remain unclear. A series of high frequency repetitive transcranial magnetic stimulation (HF-rTMS) sessions activated MEK signaling in the sensorimotor cortex, and MEK signaling was necessary for enhanced CST sprouting, regeneration and functional recovery in HF-rTMS treated SCI model mice.

## RESULTS

### Conditional, inducible gene targeting in mature cortical layer V neurons

To selectively manipulate cell-intrinsic signaling in CSNs, we utilized a *Fezf2-T2A-CreER^T2^* (*Fezf2-CreER*) knock-in mouse line (*18*) to induce expression of kinase activated (ka) B-RAF in adult CSNs. To ascertain targeting efficacy in the mature brain, *Fezf2-CreER* mice were mated with the *loxP-STOP-loxP-EgfpL10a* (*lsl-EgfpL10*) and *CAG-loxP-STOP-loxP-tdTomato* (*lsl-tdTom*) lines (*19, 20*). Tamoxifen administration to the *lsl-EgfpL10 : lsl-tdTom : Fezf2-CreER* mouse led to Cre recombination in cortical layer V neurons as indicated by EGFPL10 expression (**Fig. 1A**). Sporadic EGFPL10 positive cells were found in layer VI and hippocampus but not in the spinal cord (**Fig. 1B, C**). Bilateral injection of the anterograde tracer biotinylated dextran amines (BDA) into the motor cortex resulted in efficient and specific labeling of the CST axons (**Fig. 1D**), with the majority (83.0 ± 5.3%) of BDA labeled CST axons co-expressing tdTom (**Fig. 1E, F**). No Cre recombination was detected in the brain stem (**Fig. 1B**) or in the spinal cord grey matter (**Fig. 1C**). Injection of AAVrg-EGFP or the retrograde tracer cholera toxin subunit B (CTB) into the spinal cord of an *lsl-tdTom : Fezf2-CreER* mouse at thoracic level 8 (T8) labeled a distinct group of tdTom positive layer V neurons (**Fig. S1A**). 92.3 ± 6.2% of retrogradely CTB labeled cells in an *lsl-EgfpL10 : lsl-tdTom : Fezf2-CreER* mouse expressed EGFP, tdTom or both (**Fig. S1B-E**). Note that while tdTom filled the cell soma, dendrites (**Fig. S1D**) and axons (**Fig. 1B**), EGFPL10 was predominantly located in the cell soma, consistent with its association with ribosomes. These data demonstrate the suitability of the *Fezf2-CreER* line for selective targeting of layer V CSNs in adult mice.

**Figure 1:**
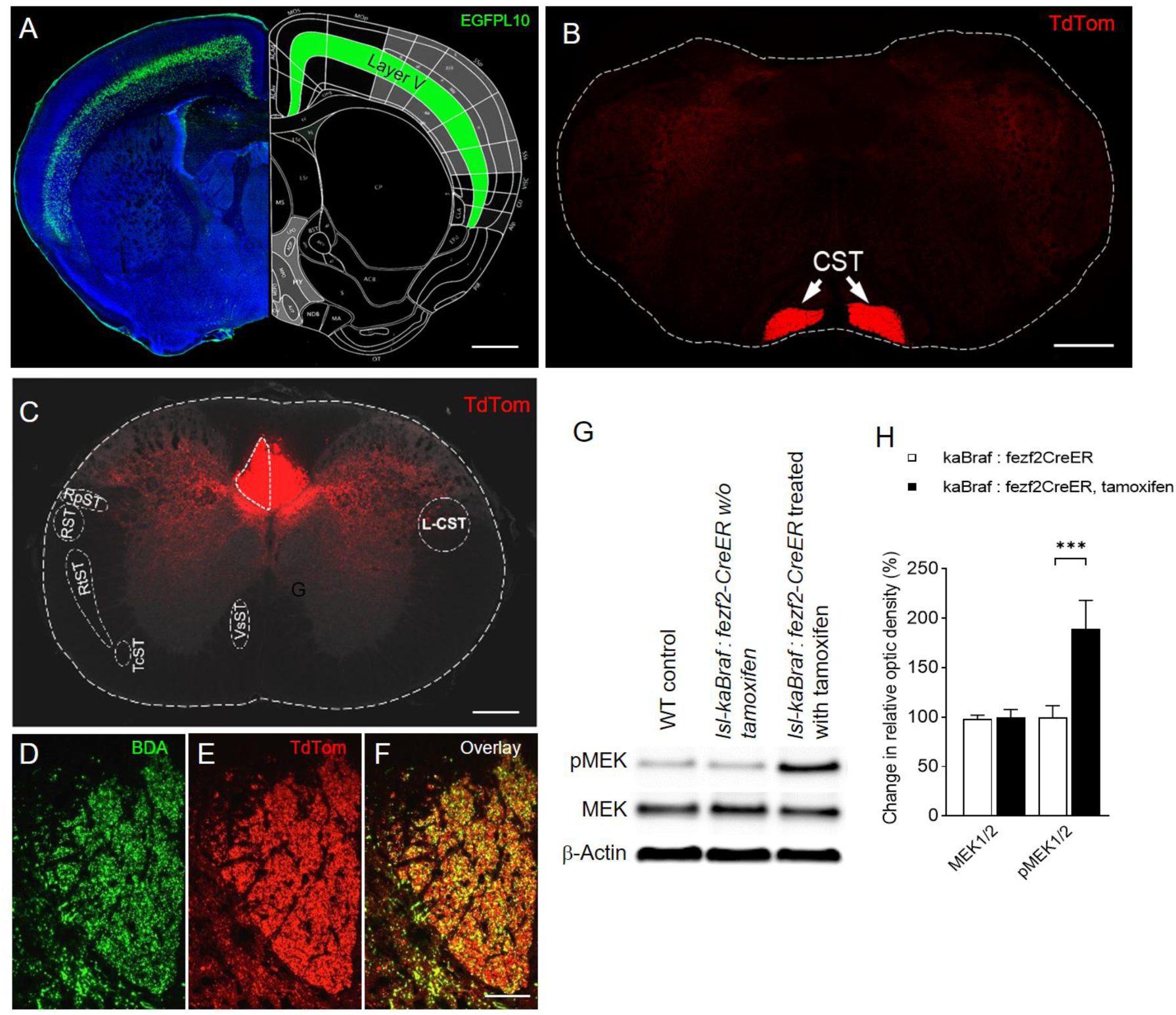
*Fezf2*-*CreER* conditionally targets layer V corticospinal projection neurons. A 6 weeks old *lsl-EgfpL10 : lsl-tdTom : Fezf2-CreER* mouse was treated with tamoxifen for 5 consecutive days to induce Cre-mediated DNA recombination; analysis was performed at age 8 weeks. (A) Left, coronal section. Green, EGFPL10; blue, Draq5. Right, reference annotation of brain regions, adapted from the Allen mouse brain atlas (Lein et al., 2007). Cortical layer V is shown in green. Scale bar, 2 mm. (B) Transverse section of medulla oblongata. Red, tdTom. Scale bar, 250 µm. (C-F) Transverse level C3 spinal cord sections. Red, tdTom. Scale bar, 250 µm. (D-F) CST of a C3 spinal cord transverse section. BDA was injected unilaterally into the motor cortex two weeks prior to fixation. (D) Green, BDA; (E) Red, tdTom; (F) Overlay of (D) and (E). Scale bar, 150 µm. (G) Western blot indicates that MEK1/2 phosphorylation is increased in the cortex of a *lsl-kaBraf : Fezf2-CreER* (ckaBraf) mouse treated with tamoxifen. (H) Densitometric quantification of (G). N = 3 per group. Error bars, mean ± SEM. **; *p* < 0.01.

Having verified the functionality of the *Fezf2-CreER* line, we bred it with mice carrying a conditional kinase activated (ka)*Braf* knock-in allele (*lsl-kaBraf*) (*10, 21*) to assess the ability of B-RAF gain-of-function to promote CST axon elongation after SCI. Expression of kaB-RAF was induced in 5 – 6 weeks old *lsl-kaBraf : lsl-tdTom : Fezf2-CreER* mice with a 5-day course of tamoxifen treatment. Cre recombination was confirmed by transcranial fluorescence imaging of tdTom. The conditional expression of kaB-RAF did not result in any overt behavioral or anatomical abnormalities in *lsl-kaBraf : lsl-tdTom : Fezf2-CreER* mice before, during or after tamoxifen administration. Tamoxifen administration increased the level of phosphorylated MEK in the cortex, indicating activation of B-RAF in the Cre expressing cells (**Fig. 1G, H**).

### Conditional B-RAF activation induces a pro-growth transcription program in CSNs

We characterized the transcriptional profile of CSNs with and without B-RAF activation by enriching the pool of actively translated mRNAs using translating ribosome affinity purification (TRAP) (*19, 22*). Intact 5-week-old *lsl-kaBraf : lsl-tdTom : lsl-Egfpl10a : fezf-CreER and lsl-tdTom : lsl-Egfpl10a : Fezf2-CreER* mice were treated with tamoxifen as above. Seven days after the last tamoxifen dose cortical tissue containing the EGFPL10 expressing cells was micro-dissected and TRAP-purified using EGFP antibodies (see Materials and Methods). The layer V neuronal markers Chst8, Etv1, Npr3, Pcp4, Vat1l, Crym, Ctip2, Foxp2 and Tle4 were enriched by 1 to 3.5 log_2_FC in the TRAP’ed samples, whereas levels of markers for layers I and II–III neurons Reln, Rasgrf2, Calb1 and Cux1 were reduced (**Fig. S2A**). Affinity purification also reduced the levels of non-neuronal cell markers (**Fig. S2B, C**). Genes with an adjusted *p* (*p*_adj_) < 0.05 were considered differentially expressed (GEO access no.: GSE201350). We verified the TRAP-seq results of select transcripts by qRT-PCR (data not shown) and *in situ* hybridization (**Fig. S3**).

A total of 2003 differentially expressed genes (DEGs) were identified, with 1047 genes up- and 956 down-regulated, including 56 up- and 80 downregulated TFs. To predict the TFs likely to be responsible for the gene expression profile we observed in kaB-RAF expressing CSNs, the full set of DEGs, excluding TFs, was uploaded to the TF enrichment analysis tools EnrichR and ChEA3 (*23, 24*). Thirty-four of the 56 upregulated TFs were predicted by both programs (**Suppl. Table S1**). The pioneer factor FOS-like 1 (FOSL1, also known as FOS-related antigen 1, FRA-1) topped the list of 200 potentially relevant TFs predicted by ChEA3, with a score of 17.8. Notably, several basic leucine zipper (bZIP) family TFs were among the top upregulated TFs (**Fig. 2A**): FOSL1 and cAMP-responsive element binding protein 3-like 1 (CREB3L1) were increased by 4.5 and 2.9 log_2_-fold change (log_2_FC), respectively, and nuclear factor erythroid 2-like factor 3 (NFE2L3), CCAAT enhancer binding protein *δ* (CEBPD), melanocyte-inducing transcription factor (MITF) and v-maf musculoaponeurotic fibrosarcoma oncogene family protein F (MAFF) were up by 1.78, 1.09, 1.06 and 0.92 log_2_FC, respectively. The mRNA level of JUN, a well-established hub regulator in PNS axon growth and regeneration (*25–28*) was high at baseline compared to other bZIP TFs, and further increased by 0.45 log_2_FC in kaB-RAF expressing CSNs. Interestingly, the TFs upregulated by B-RAF gain-of-function overlapped substantially with the set of TFs that are upregulated in the beginning phase of axon regeneration in axotomized zebrafish retinal ganglion cells (RGCs) (*29*): six of the TFs upregulated by B-RAF are orthologs of 9 out of the set of 11 zebrafish TFs induced in regeneration-competent zebrafish RGCs within 2 days of axotomy (**Fig. 2C**). Moreover, three of the B-RAF-induced TFs, FOSL1, WT1 and EGR2 have been recognized as pioneer factors modulating chromatin accessibility to initiate regulatory events (*30–34*). FOSL1 and WT1 were also upregulated in zebrafish RGCs, rat dorsal root ganglion (DRG) sensory neurons and mouse facial motor neurons soon after nerve injury (*28, 29, 35*). TFs downregulated by B-RAF activation include Krüppel-like factor-4 (KLF4, log_2_FC = -0.83), which is also a pioneer factor and known to limit axon regenerative capacity in RGCs (*36*) and CSNs (*37*). Finally, among the 56 TFs upregulated by B-RAF gain-of-function, FOSL1, CEBPD, JUN, EGR2, ELK3, CBFB, STAT3/4/6 and ETV4 are considered to be regeneration-associated genes (RAGs) in CNS or PNS neurons (*25, 28, 38*). Collectively, these data support our hypothesis that conditional activation of B-RAF signaling induces a pro-growth transcription program in mature corticospinal neurons.

**Figure 2:**
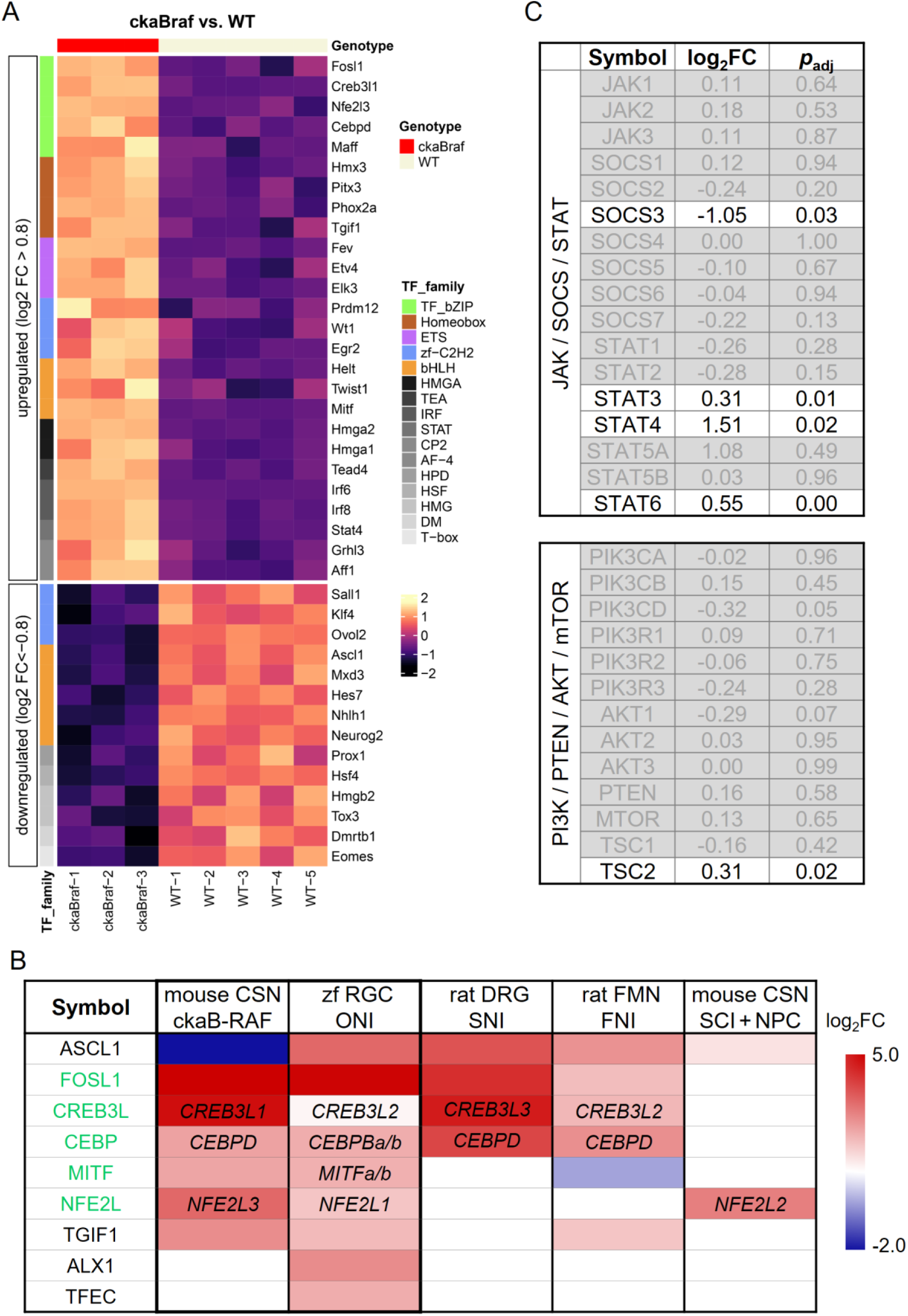
Translatome analysis of DEGs induced by B-RAF activation in CSNs. (A) Heatmaps of DE-TFs in FEZF2+ neurons induced by B-RAF GOF. Top up- and downregulated TFs (log_2_FC > 0.8 or log_2_FC < -0.8) are listed. TFs are clustered based on structural family. Color scale indicates change in expression level (z-score, *p_adj_* < 0.05). (B) Comparison of TFs induced by B-RAF activation with TFs upregulated in regeneration-competent zebrafish retinal ganglion cells (zf RGC) and other published CNS and PNS axon regeneration studies. Six of the 9 TF isoforms identified as early reprogramming factors in regenerating zf RGC (*29*) were upregulated by B-RAF activation in CSNs. CSN/ckaB-RAF, this study; zf RGC/ONI, zf RGCs 2 days post optic nerve injury (*29*); rat DRG/SNI, DRG neurons 7 days post sciatic nerve injury (*35*); facial MN/FNI, facial motor neurons 1 and 4 days after facial nerve injury (*28*); CSN/SCI + graft, CSNs 3 and 7 days post SCI growing into a neural precursor cell graft (*47*). SNI, sciatic nerve injury; FMN, facial motor neurons, SCI, spinal cord injury. Transcripts were included if their up- or downregulation met the significance threshold of *p*_adj_ < 0.05 for the CSN/ckaB-RAF, zf RGC/ONI, facial MN/FNI and CSN/SCI + NPC graft data, or *p* < 0.05 for the rat DRG/SNI data. Gene symbols are spelled out in cells when the upregulated genes are different but belong to the same paralogous group. Two zf TFs (CEBPBa/b, MITFa/b) have isoform duplicates (*89*). Color indicates the one with reported higher change in expression. (C) Expression of key regulatory genes in JAK/SOCS/STAT and PI3/PTEN/AKT/mTOR pathways. SOCS3 transcript was reduced and STAT3/4/6 transcripts were elevated in CSNs expressing kaB-RAF. Only one gene in the PI3K/mTOR pathway, TSC2, was modestly upregulated. Genes with a *p_ad_*_j_ > 0.05 and genes downregulated by B-RAF activation are shown greyed out.

### B-RAF activation elicits SOCS – STAT, but not PI3K – mTOR signaling

Interleukin-6 family cytokines and cytokine-activated JAK – STAT signaling have been implicated in PNS and CNS axon regeneration (*39–42*). B-RAF activation downregulated the JAK/STAT antagonist SOCS3 (log_2_ FC = -1.05) and upregulated the TFs STAT3/4/6 (log_2_ FC = 0.31, 1.5, 0.55, respectively), indicating that B-RAF activation increased JAK/STAT signaling (**Fig. 2B**). In contrast, B-RAF activation did not directly regulate the expression of mRNAs related to the PTEN – mTOR pathway (except for TSC2, log_2_FC = 0.3) in CSNs, another pathway strongly implicated in multiple CNS regeneration paradigms (*43–46*). TRAP-seq analysis of CSNs from two PTEN conditional knockout mice (*Pten ^f/f^ : lsl-tdTom : lsl-Egfpl10a : fezf-CreER*) revealed a distinct gene expression profile from conditional B-RAF activation (**Fig. S4**). Gene ontology analysis indicates that kaB-RAF expression activates transcripts involved in multiple signaling pathways related to axon extension (**Table S2**). Comparing Ingenuity Pathway Analysis (IPA) results from our dataset with previously published data from regenerating sensory dorsal root ganglion (DRG) neurons, facial motor neurons and CSNs (*28, 35, 47*) uncovered pathways that appear to be shared among all four experimental paradigms, specifically CREB signaling, GPCR signaling and focal adhesion kinase signaling (**Table S2**). In contrast, the mTOR pathway was activated only in regenerating facial motor and corticospinal neurons, and neuroinflammation pathways were activated in all injured and regenerating neuron types, but not in the kaB-RAF expressing CSNs, which were intact.

### B-RAF activation facilitates contralateral sprouting and *de novo* synaptogenesis after unilateral pyramidotomy

Human spinal cord injuries are usually incomplete. Spontaneous sprouting of collaterals from spared axons is hypothesized to underlie observed functional restoration in human patients as well as rodent SCI models (*48–50*). The actual capacity of spontaneous axon collateral sprouting in adult CNS is very limited, however, so interventions to increase collateral sprouting could enhance functional recovery. We utilized the unilateral pyramidotomy model (uPx) to assess the effects of B-RAF activation on sprouting of spared CST axons. uPx led to complete denervation of the contralateral CST (**Fig. S5**). *lsl-kaBraf : lsl-tdTom : fezf-CreER* mice were treated with tamoxifen prior to uPx. In contrast to dorsal hemisection (dHx), where severed axons can grow across the lesion site to establish new connections, no axon regeneration is possible after uPx. Instead, collateral branches from contralateral intact axons sprout across the midline to establish new connectivity (*50*). Within 4 weeks following uPx, we observed approximately 5-fold enhanced contralateral sprouting in *lsl-kaBraf : lsl-tdTom : Fezf2-CreER* mice, compared to the very limited contralateral axon projections seen in control littermates lacking kaB-RAF expression in the CSNs (**Fig. 3A-E**). We then asked whether the new collaterals are capable of forming synaptic connections with resident neurons in the denervated CST. Mice subjected to uPx, with or without induction of kaB-RAF expression, received stereotactic injections of anterograde trans-synaptic AAV–WGACreER (**Video S1**) into motor cortex contralateral to the injury 2 weeks post surgery (**Fig. 3F**). As expected, very few tdTom positive cells were detectable in the gray matter of the denervated side of the CST in control mice without B-RAF activation (**Fig. 3G**). In contrast, mice expressing kaB-RAF displayed a ∼6 fold higher number of labeled interneurons in the denervated CST (**Fig. 3H, I**), indicating *de novo* synapse formation fromcollateral axon branches originating from the intact contralateral CST.

**Figure 3:**
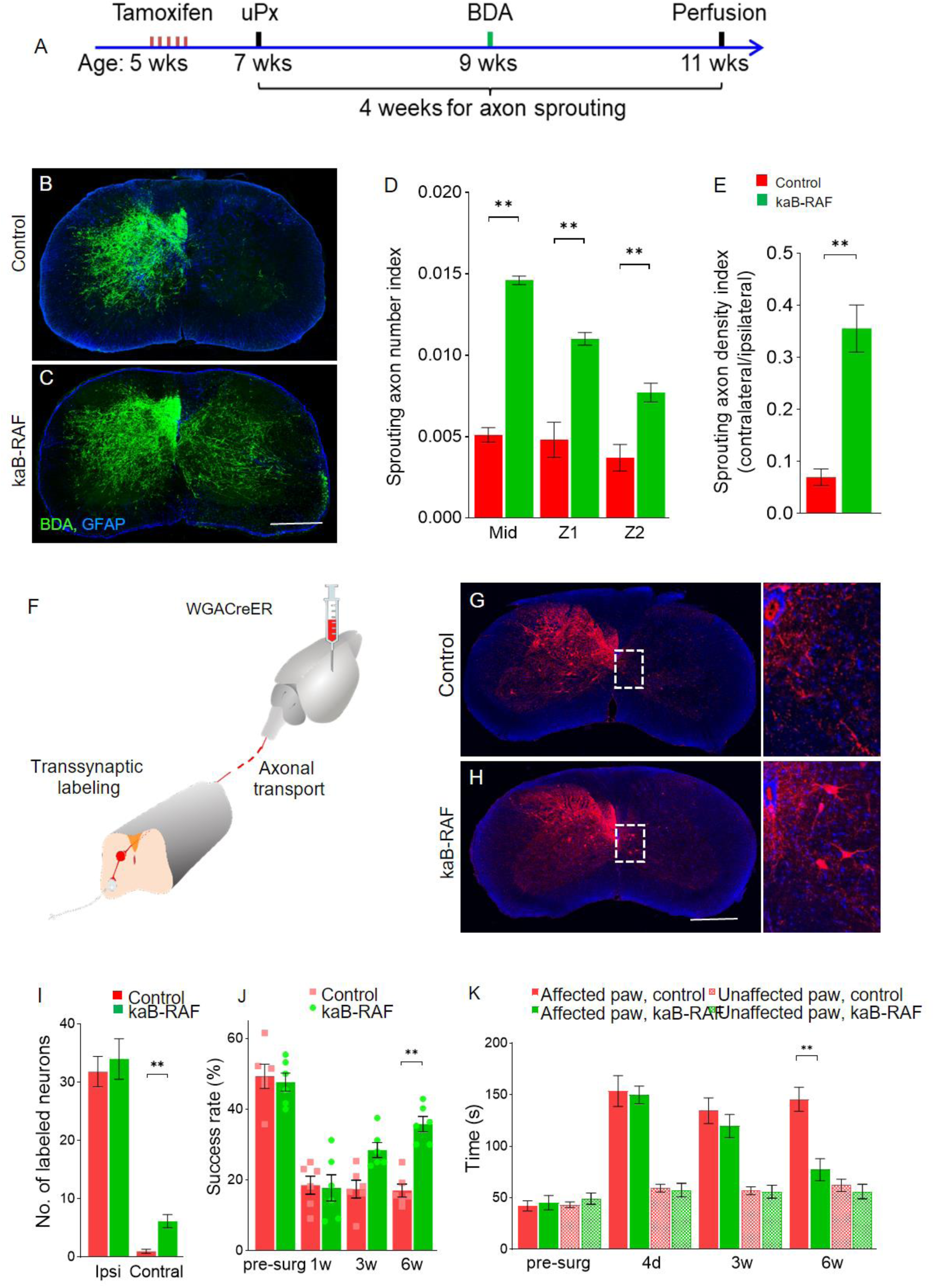
kaB-RAF promotes CST axon sprouting, synaptogenesis and recovery of skilled forelimb function after unilateral pyramidotomy (uPx). (A) Experimental timeline. (B, C) Transverse sections of level C5 spinal cord from tamoxifen treated (B) *lsl-tdTom : Fezf2-CreER* and (C) *lsl-kaBraf : lsl-tdTom : Fezf2-CreER* mice 6 weeks after uPx. Green, BDA; blue, GFAP; scale bar, 200 μm. (D, E) Quantification of BDA labeled sprouting axons crossing the midline. (D) Axon number index at midline (Mid) and extending 200 µm (Z1) and 400 µm (Z2) beyond the midline. (E) Axon density index. \*\**p* < 0.001, two-way ANOVA with Bonferroni *post-hoc* correction. N = 5 per group. Error bars, mean ± SEM. (F–H) New synaptic connections following uPx. (F) Four weeks after uPx surgery, an AAV expressing a transsynaptically WGA-CreER was injected into the motor cortex contralateral to the lesion side. Transverse sections of C7 spinal cord are shown from adult (G) *lsl-tdTom : Fezf2-*CreER and (H) *lsl-kaBraf : lsl-tdTom : Fezf2-CreER* mice 10 days after AAV injection. Red (tdTom) labels local propriospinal interneurons contacted by newly sprouted axon collaterals. Blue, GFAP. (I) Quantification of transsynaptically labeled interneurons in gray matter. One-way ANOVA with Bonferroni *post-hoc* correction, \*\**p* < 0.001. N = 5 per group. Error bars, mean ± SEM. (J) Pellet grasping assay. Mice conditionally expressing kaB-RAF in corticospinal neurons recover significant pellet grasping ability following uPx. \*\**p* < 0.05. (K) Patch removal test. The time between noticing and the successful removal were measured. \*\**p* < 0.001. N = 5 per group. Error bars, mean ± SEM.

### B-RAF activation improves recovery of forelimb motor control after unilateral pyramidotomy

To determine the contribution of B-RAF mediated axon contralateral sprouting to the recovery of motor control in mice subjected to uPx, we quantified voluntary motor performance using a single pellet grasping assay (**Fig. 3J**) (*51, 52*) and a patch removal test (**Fig. 3K**) (*53, 54*). Following tamoxifen treatment, *lsl-kaBraf : lsl-tdTom : Fezf2-CreER* mice and control littermates were trained daily for 2 weeks in the grasping paradigm. The preferred paw for each mouse was determined during training, and uPx was then performed on the contralateral side (to disable the preferred paw). Prior to uPx, all mice displayed similar pellet grasping success rates, 49.3 ± 3.4% for *lsl-kaBraf : lsl-tdTom : Fezf2-CreER* and 47.6 ± 2.6% for *lsl-tdTom : Fezf2-CreER.* Following uPx surgery, success rates dropped to the same extent in both groups (18.5 ± 2.6% vs 17.7 ± 3.7%). Over time, the performance of mice expressing kaB-RAF gradually improved. Six weeks after surgery, their success rate reached 35.8 ± 2.1%, twice as successful as the control group (17.0 ± 1.8%). In the patch removal test, a measure of dexterity, mice subjected to uPx and expressing kaB-RAF recovered to almost normal level by 6 weeks after the surgery, while littermates lacking kaB-RAF did not improve significantly (**Fig. 3K**). Taken together, B-RAF-mediated enhancement of axon collateral sprouting correlated with increased *de novo* synaptogenesis in the denervated CST, and with improved recovery of skilled forelimb function.

### B-RAF activation promotes corticospinal axon regeneration after dorsal hemisection

To test the potency of B-RAF activation as a way to promote CNS axon regeneration, we subjected the mice to dorsal hemisection (dHx) injury (*44, 55*). *lsl-kaBraf : lsl-tdTom : fezf-CreER* mice were treated with tamoxifen for 5 days to induce kaB-RAF expression, rested for 7 days and then underwent dHx surgery at level T8. We allowed 6 weeks for the axons to regenerate (**Fig. 4A**). BDA was injected into the motor cortex to anterogradely label descending CST axons two weeks before the end of the experiment; injection coordinates were determined by retrograde labeling of T8 corticospinal projection neurons (**Fig. S6**). Robust CST axon outgrowth beyond the lesion epicenter was evident in conditional B-RAF GOF mice 6 weeks post dHx (**Fig. 4B-D**). To confirm the cortical origin of these axons, AAVrg-EGFP was unilaterally injected into the spinal cord gray matter 2 mm caudal to the lesion epicenter. EGFP positive cells were detected in layer V tdTom positive cells of the *lsl-kaBraf : lsl-tdTom : Fezf2-CreER* mouse, but not in the control (**Fig. 4E**). In addition, all regenerating axons caudal to the lesion epicenter were tdTom positive, indicating successful Cre recombination and therefore kaB-RAF expression in the regenerating CSNs (**Fig. S6G**).

**Figure 4:**
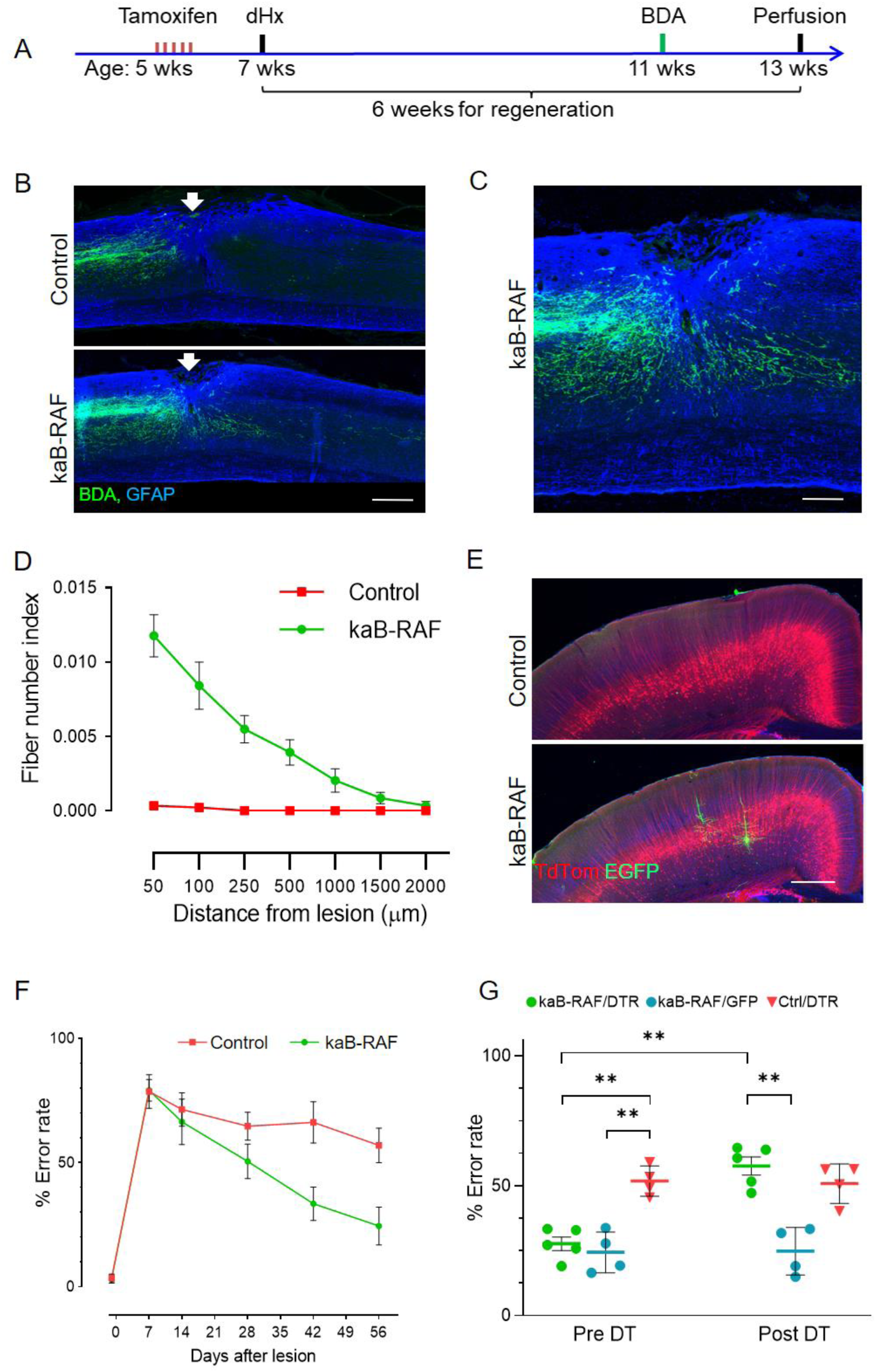
Conditional B-RAF activation promotes axon regeneration after a spinal cord T8 dorsal hemisection (dHx) injury. (A) Experimental timeline. KaB-RAF expression was induced by tamoxifen administration in 5 weeks old *lsl-kaBraf : lsl-tdTom : Fezf2-CreER* mice. dHx was performed two weeks later, and axon regeneration analyzed 6 weeks post injury. (B, C) Representative images of sagittal sections from the lesion site 8 weeks after T8 dHx. Green, anterograde tracer BDA; blue, GFAP. Arrows indicate lesion epicenters. Scale bar, 250 µm. (C) high magnification image of (B). (D) Quantification of labeled axons in the spinal cord caudal to the lesion site after dHx. *p* < 0.0001, two-way ANOVA followed by Bonferroni *post-hoc* correction. Error bars, mean ± SEM. N = 6 per group. (E) Regenerating CSNs labeled by AAVretrog-EGFP injected into spinal cord gray matter 2 mm caudal to the lesion epicenter. Scale bar, 200 µm. (F) Improved functional recovery after a T8 dHx. Mice were trained to cross a horizontal ladder with irregularly spaced rungs. Stepping error rates of kaB-RAF expressing and control mice were quantified. Error bars, mean ± SEM. Two-way ANOVA followed by Bonferroni *post-hoc* correction, *p* < 0.0001. N = 6 for each group. (G) Ablation of the CSNs extending axons across the lesion center reverses the functional recovery of B-RAF GOF mice. Diphtheria toxin receptor (DTR) was conditionally expressed in CreER positive CSNs by retrograde gene transfer and subsequent tamoxifen induced recombination. Mice were re-tested on the ladder 2 weeks after diphtheria toxin (DT) administration. Error bars, mean ± SEM. Two-way ANOVA followed by Bonferroni *post-hoc* correction, \*\**p* < 0.001. N = 4 or 5 per group.

To ensure the completeness of the CST lesion, transverse sections 5 mm caudal and rostral to the lesion epicenter were checked for spared axons (**Fig. S6**). No BDA staining was detected in sections caudal to the lesion epicenter (**Fig. S6C, D**). Furthermore, PKCγ, a marker of CST axons, was completely abolished distal to the lesion (**Fig. S6E, F**). No BDA labeled axons were detected distal to the lesion site in control *lsl-tdTom : Fezf2-CreER* mice.

### Recovery of skilled stepping after dorsal hemisection, dependent on CST axon regeneration

Recovery of CST-dependent skilled motor function was assessed on the horizontal ladder with irregularly spaced rungs (*56*). Prior to surgery, tamoxifen treated control *lsl-tdTom : Fezf2-CreER* and kaB-RAF expressing *lsl-kaBraf : lsl-tdTom : Fezf2-CreER* mice displayed similar low stepping error rates (3.5 ± 0.5%, mean ± SEM). The error rates of both groups dramatically increased to 78.7 ± 2.2% for the control and 79.1 ± 1.8% for kaB-RAF mice one week post surgery. Beginning at 2 weeks post dHx, mice expressing kaB-RAF in CSNs started to outperform the control group. The difference became statistically significant 4 weeks after surgery. At 8 weeks post surgery, the median error rate of kaB-RAF expressing mice was 24.4 ± 3.1%, compared to 56.9 ± 2.2% in the controls (**Fig. 4F**). To ascertain that CST axon regeneration was indeed the cause of skilled locomotion recovery, we ablated the CSNs that had regenerated axons beyond the T8 dHx lesion site using a viral retrograde targeting approach (*57*). A lentivirus (LV) vector harboring a conditional diphtheria toxin receptor (DTR) expression cassette (HiRet-FLEX-DTR, *58*) was bilaterally injected into the spinal cord gray matter 2 mm caudal to the lesion epicenter, followed by tamoxifen administration to trigger DTR expression in CreER-expressing CSNs that incorporated the LV. Five days after the last tamoxifen treatment, diphtheria toxin (DT) was administered to specifically ablate cells expressing DTR. The horizontal ladder walking assay was repeated one week after the lentivirus injection and 2 weeks after DT administration. Error rates remained essentially unchanged after LV injection: 27.6 ± 2.6% for mice expressing kaB-RAF and 57.6 ± 3.5% for the control group (**Fig. 4G**). Following DT administration, the error rates of kaB-RAF expressing mice rose to 51.8 ± 3.9%, matching the error rate of the control group (50.8 ± 3.8%), which remained unaffected by DT treatment. In a control dHx group expressing kaB-RAF and injected with a lentivirus expressing HiRet-GFP, DT administration had no effect on the error rate (24.3 ± 4.0% vs 24.8 ± 4.6%) (**Fig. 4G**). These data indicate that the kaB-RAF-dependent growth of CST axons across the lesion site causally contributes to the recovery of voluntary motor control after dHx.

### High frequency rTMS activates MEK1/2 in sensorimotor cortex

Membrane depolarization *in vitro* can directly activate MEK1/2 via calcium influx (*59, 60*), and high frequency electrical stimulation activates RAS – RAF signaling in CA1 pyramidal neurons (*61*). Here, we treated awake mice with a course of HF-rTMS to determine its efficacy in activating RAF – MEK signaling *in vivo*. Mice received one HF-rTMS session daily, with 75 pulses at 15 Hz per train, 10 trains per session, delivered at 120% of resting motor threshold (RMT) intensity. Simulation of the electric field (E-field) distribution (*62*) induced by our HF-rTMS protocol shows that the expected diffuse ring-shaped stimulation pattern (*63*) encompasses the whole mouse brain. The calculated peak amplitudes of E-fields in the forebrain range between 40-60 V/m (**Fig. 5A**), which is in line with the 35 or 45 V/m that are typically employed to depolarize or hyperpolarize CNS neurons in rodents or non-human primates *in vivo* (*64, 65*). Since no antibody is available to directly assess the activation level of B-RAF, we quantified the phosphorylation of its canonical effector MEK1/2 as a measure of B-RAF activity. Daily HF-rTMS treatment over 5 consecutive days increased the level of pMEK1/2 in the cortex of wild type mice to 224 ± 41% of that in sham treated mice (**Fig. 5B, C**), indicating activation of MEK signaling. Stimulation at 50% RMT instead of 120% elicited no hindlimb muscle twitch, and no increase in pMEK1/2 after a 5-day treatment course (**Fig. S7**). We did not observe any spasticity or other abnormal behaviors in HF-rTMS treated mice during the entire experimental period.

**Figure 5:**
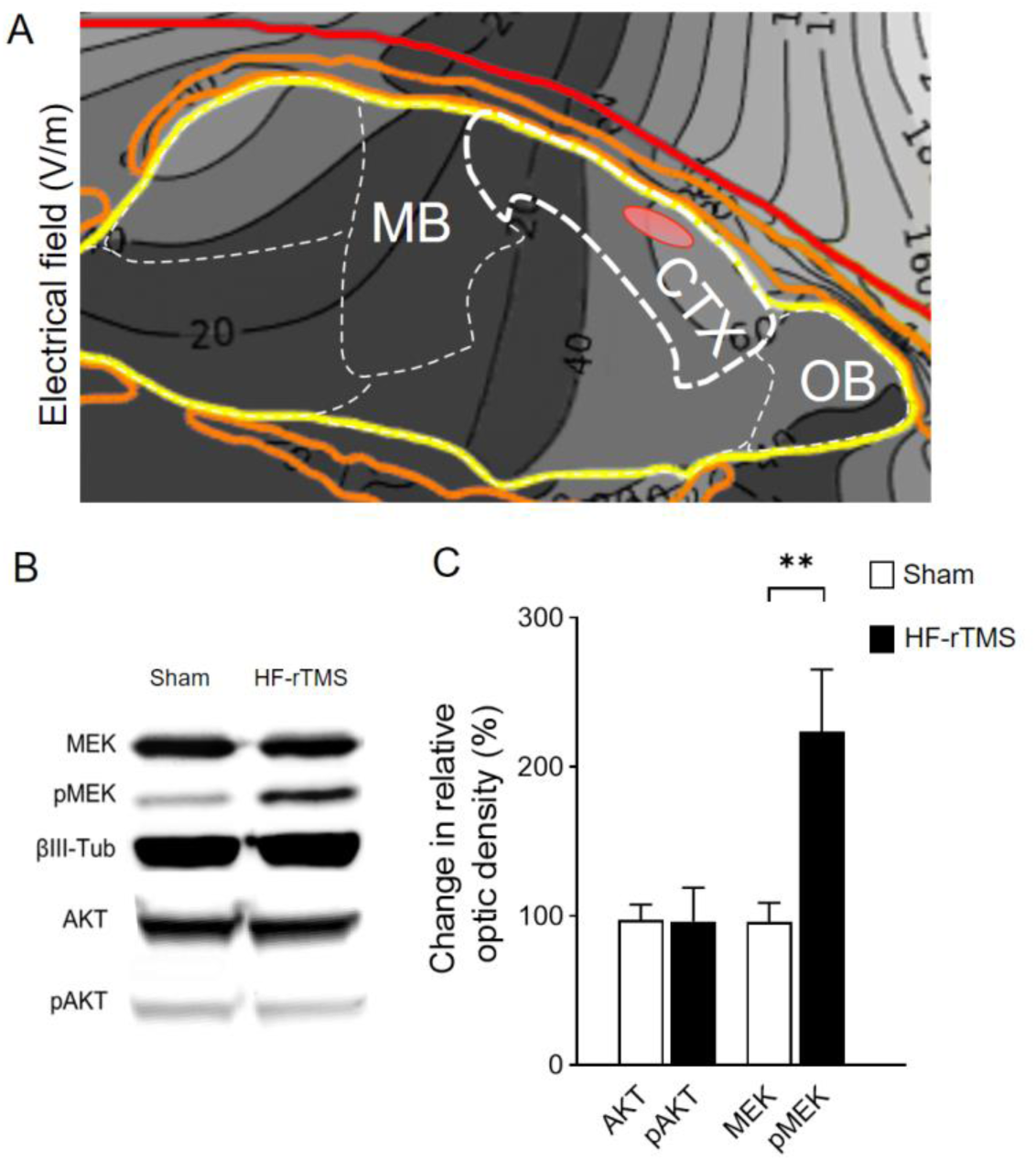
HF-rTMS activates MAP kinase signaling in the cortex of adult wild type mice. (A) Boundary-element simulation of the TMS-induced E-field in a generic mouse head by the Cool-40 rat coil at 35% of maximum stimulator output (MSO). (B, C) Increased MEK1/2 phosphorylation in cortex following cranial HF-rTMS. Red oval indicates the approximate location of corticospinal neurons projecting to spinal cord level T8. MB, midbrain; CTX, cortex; OB, olfactory bulb. (B) Mice were treated with HF-rTMS (15 Hz) at 120% of their RMT (approximately 35% MSO) daily for 5 consecutive days. Representative Western blot showing elevated pMEK1/2 levels in HF-rTMS treated cortex. No change in pAKT was detected. (C) Densitometric quantification of AKT and MEK total protein and protein phosphorylation. ** *P* < 0.001. ANOVA with Bonferroni correction. N = 3 per group. Error bars, mean ± SEM.

### HF-rTMS promotes CST axon sprouting, regeneration and functional recovery after CST injury

We then assessed the potency of HF-rTMS in enhancing CST axon sprouting. Wild type mice aged 5 – 6 weeks were treated with tamoxifen (to maintain comparable conditions to the conditional MEK1/2 ablation experiment below), followed by uPx surgery. Starting one day post surgery, HF-rTMS was administered to awake mice, daily for 4 weeks. HF-rTMS treatment increased sprouting of collateral branches across the midline from the intact into the denervated side of the spinal cord (**Fig. 6A-E, H**). We quantified voluntary motor performance with the single pellet grasping assay using the same protocol described above. Prior to uPx, all mice displayed similar pellet grasping success rates, and success rates dropped to the same extent in both groups after uPx surgery. HF-rTMS was administered as above. Over time, the performance of HF-rTMS treated mice improved to a greater extent than that of sham controls. Six weeks post surgery, their success rate reached 36.4 ± 5.1 % (mean ± SEM) whereas the rate of the control group was 20.5 ± 3.5% (**Fig. 6I**).

**Figure 6:**
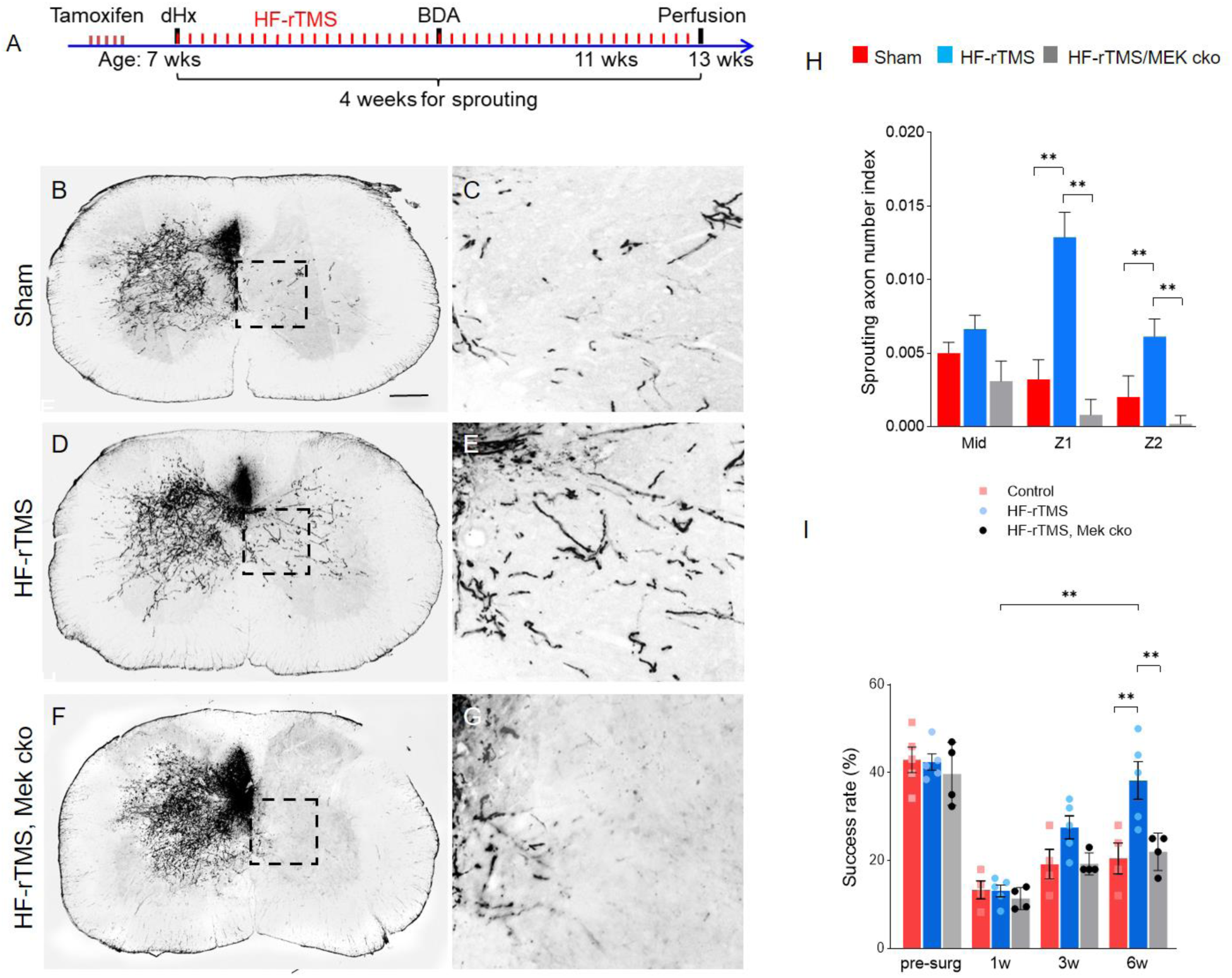
HF-rTMS increases CST axon sprouting after unilateral pyramidotomy, dependent on MEK1/2 signaling. (A) Experiment timeline. (B – G) Spinal cord level C5 transverse sections 8 weeks after uPx. rTMS was administered daily starting 2 days after injury. Axons were labeled by unilateral BDA injection. (A, B) Baseline level of contralateral sprouting in a sham-treated mouse. (C, D) HF-rTMS-induced contralateral sprouting. (E, F) Genetic ablation of RAF effectors blocked rTMS induced axon sprouting in *lsl-Mek1^f/f^ : Mek2^-/-^ : Fezf2-CreER ^T2^* mice treated with tamoxifen. Scale bar, 50 µm. (H) Quantification. N = 4 per group. ** *P* < 0.001, one-way ANOVA with Bonferroni correction. (I) Pellet grasping assay. HF-rTMS treatment enhanced recovery of pellet grasping ability following uPx only in mice with intact endogenous MEK1/2. N = 4 per group. ** *P* < 0.05.

We then applied HF-rTMS to mice subject to dHx surgery. Wild type mice at age 5 – 6 weeks were treated with tamoxifen, followed by dHx surgery. Starting one day post surgery, HF-rTMS was administered to awake mice, daily for 6 weeks. This procedure triggered significant CST axon growth across the lesion site (**Fig. 7A-C, E**). rTMS–treated mice also recovered better stepping ability on the horizontal ladder compared to sham-treated controls (**Fig. 7F**).

**Figure 7:**
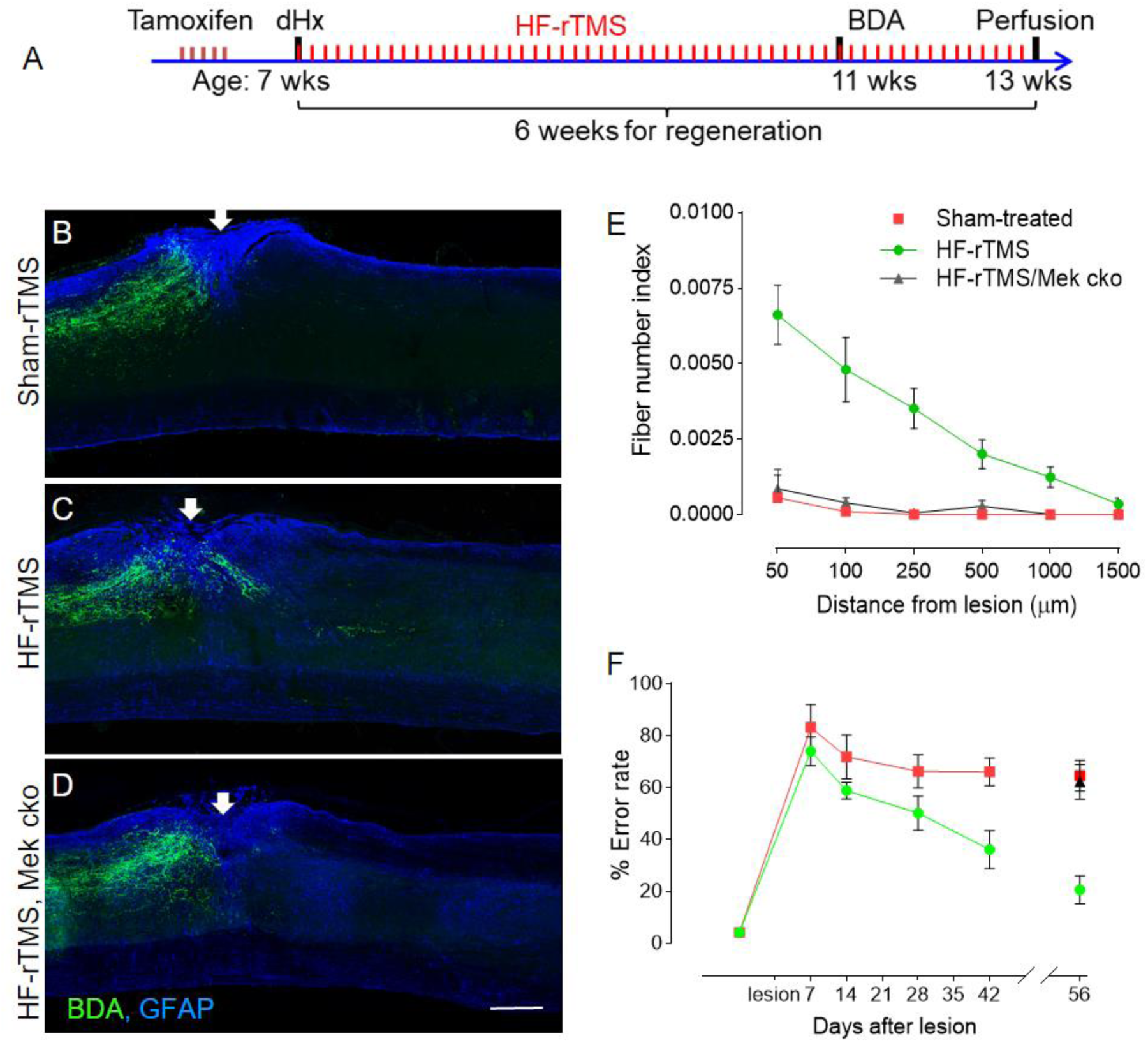
HF-rTMS facilitates CST axon regeneration and functional recovery after T8 dorsal hemisection, depending on activity of Mek1/2. (A) Experimental timeline. (B – D) Representative images of spinal cord sagittal sections 10 weeks after dHx from (B) wild type sham-treated, (C) wild type HF-rTMS treated, and (D) HF-rTMS treated *Mek1^f/f^ : Mek2^-/-^ : Fezf2-CreER* mice. Scale bar, 300 µm. (E) Quantification of axon regeneration. *P* < 0.0001, two-way ANOVA with Bonferroni correction. N = 6 per group. Error bars, mean ± SEM. (F) Recovery of skilled stepping in HF-rTMS treated mice. N = 6 per group. *P* < 0.001, two-way ANOVA with Bonferroni correction. Error bars, mean ± SEM.

### HF-rTMS-induced CST axon sprouting and regeneration depends on endogenous MEK signaling

Loss of MEK1/2 is sufficient to block B-RAF induced axon growth in the optic nerve after injury (*10*). To determine the involvement of MEK signaling in HF-rTMS induced axon growth, we conditionally deleted the kinases MEK1 and MEK2 in CSNs. *Mek1 ^f/f^ : Mek2 ^-/-^ : Fezf2-CreER* mice were treated with tamoxifen and subjected to uPx surgery, then underwent daily HF-rTMS treatment as above. Conditional ablation of MEK1/2 completely blocked HF-rTMS–induced CST axon sprouting (**Fig. 6F, G, H**). Consequently, post-injury recovery of fine motor control was reduced in comparison to HF-rTMS treated mice carrying intact Mek1/2 alleles (**Fig. 6I**). Similar results were observed in *Mek1 ^f/f^ : Mek2 ^-/-^ : Fezf2-CreER* mice subject to dHx surgery: HF-rTMS was unable to promote axon regeneration (**Fig. 7D, E**) or to improve post-SCI stepping recovery in these mice (**Fig. 7F**). *Mek1 ^f/f^ : Mek2 ^-/-^ : Fezf2-CreER* mice displayed no obvious abnormalities before or after tamoxifen treatment. We have previously shown that loss of MEK1/2 in RGCs has no effect on RGC survival (*10*) and also did not observe any cortical neuron loss in *Mek1 ^f/f^ : Mek2 ^-/-^ : Fezf2-CreER* mice (data not shown).

We conclude that HF-rTMS activates MEK in the cortex, and that MEK activation in CSNs is necessary for and correlates with substantial CST axon contralateral sprouting or regeneration post injury, as well as with improved motor recovery. Collectively, these findings establish a crucial role for MEK-dependent signaling in mature CSN axon growth and sprouting.

## DISCUSSION

Our data show that activation of B-RAF–MEK signaling in corticospinal neurons (CSNs), by genetic means or by HF-rTMS, promotes robust elongation and sprouting of CST axons in mouse models of spinal cord injury, correlated with improved fine motor control. TRAP-seq analysis revealed that B-RAF activation in mature CSNs elicits expression of a set of TFs that overlaps substantially with early response TFs induced in regeneration-competent zebrafish RGCs (*29*). There is a conspicuous induction of bZIP family TFs in both systems, including the presumptive pioneer factor FOSL1 which can dimerize with JUN to form AP-1 complexes and pioneers chromatin access (*30, 66*). JUN is recognized as a hub regulator of PNS axon regeneration (*25, 28*). Peripheral axotomy-induces the expression of JUN in sensory neurons, as well as a subset of its dimerization partners that predominantly bind distant enhancer elements rather than proximal promoters (*27*). Conditional deletion of JUN diminishes axon regeneration in facial motor neurons (*67, 68*). Overexpression of JUN promotes neurite elongation in DRG neurons but not in cortical neurons (*69*), The two JUN partners induced by B-RAF activation in CSNs, FOSL1 and CREB3L1, have not yet been characterized as to their function in the cortex, but are likely part of a molecular executive program that underlies B-RAF mediated axon elongation. FOSL1 has been reported to suppress KLF4 expression in embryonic stem cells (*30*). KLF4 antagonizes STAT3 and thereby limits JAK – STAT3 driven axon regeneration of adult mouse RGCs (*70*). Our finding that B-RAF activation increases FOSL1 and STAT3 transcripts while reducing KLF4 potentially links B-RAF to JAK – STAT3 signaling in CSNs.

At an intermediate stage of regeneration, when axons begin their target innervation, injured zebrafish RGCs upregulate a different set of TF_S_ that include the pioneer factor WT1, as well as genes in the PTEN–mTOR pathway (*29*). B-RAF activation induced WT1 expression in CSNs, but did not significantly alter the expression of PTEN–mTOR signaling mediators. Indeed, conditional Pten deletion in adult FEZF2+ neurons resulted in a TF expression profile very different from that induced by B-RAF activation. Thus, B-RAF and mTOR are likely to mobilize distinct genetic programs that may act synergistically to drive axon regenerative growth as we had previously reported for the injured optic nerve (*10*).

Neuronal precursor cells (NPC) grafted into the injured spinal cord can promote axon regeneration of CSNs (*47, 71*). Notably, JUN and JUN-associated bZIP TFs induced by B-RAF and zebrafish RGC axotomy were not upregulated either 3 days or 2 weeks after SCI, with or without NPC grafting, with the exception of NFE2L2. It appears that the two experiments, although both promote CSN axon regeneration, recruit distinct genetic programs, a conclusion that is also supported by gene ontology analysis (**Table S2**). Finally, comparing the translatome induced by B-RAF activation in CSNs with that induced in intestinal epithelial cells (IECs) where B-RAF activation is oncogenic (GSE106330, *72*), we found minimal overlap with none of the three pioneer factors, FOSL1, WT1 or EGR2, induced in IECs, demonstrating that B-RAF signaling regulates vastly different genetic programs and biological functions depending on cellular context.

B-RAF-mediated axon regeneration after dHx was robust, but we were not able to detect newly connected interneurons via transsynaptic tracing. Nevertheless, the DTR/DT ablation approach provides indirect evidence of axon reconnection-mediated functional recovery following dHx. The unilateral pyramidotomy (uPx) model of SCI, where axon regeneration into the spinal cord is not possible, allows the specific exploration of B-RAF function in axonal sprouting and synaptogenesis. Sprouting of collaterals from spared axons is of great clinical relevance as the majority of human spinal cord injuries are incomplete, with spared (intact) axons remaining in various spinal tracts (*73*). Newly sprouted collaterals from spared axons can be recruited into new circuitry and contribute to functional recovery in pre-clinical SCI models (*50, 74–76*), and particularly in primates (*77*). Sprouting of new connections from spared fibers is also likely the foundation for new interventions that enable paraplegic patients to recover walking ability (*78*). To identify synaptic connections from newly sprouted CST collaterals, we documented the transport of a transsynaptic tracer from CSNs into postsynaptic spinal interneurons. The exact phenotype of the post-synaptic neurons as well as the electrical functionality of the new connections remain to be investigated in future studies. Jayaprakash et al. (*79*) developed an optogenetic methodology to demonstrate *bona fide* synaptic transmission arising in the uPx paradigm, using viral SOX11 overexpression to drive collateral sprouting into the denervated CST. This group did not observe functional recovery in the pellet grasping or horizontal ladder assays (*80*), while we did find improved performance. Possible explanations might involve higher numbers of new synapses with activation of B-RAF, or differences in the exact subpopulations of postsynaptic spinal neurons participating in synaptogenesis.

Finally, we found that daily HF-rTMS treatments enabled robust CST axon regeneration after dHx as well as contralateral sprouting following uPx. HF-rTMS elevated the level of pMEK, and as expected, its effect promoting axon elongation was strictly dependent on the endogenous MEK1/2. While we did not detect significant phosphorylation/activation of AKT by HF-rTMS in our experiments, HF-rTMS administered at 400Hz can activate both the MAP kinase and mTOR–S6 kinase pathways in cortical neurons (*81*). HF-rTMS has also been shown to increase the expression of BDNF and other growth factors in rat and mouse brains (*82–85*) and increases glucose uptake in deep layer pyramidal neurons (Cermak et al., 2020), all of which may collectively contribute to the regenerative growth of CST axons we observed. The mechanisms by which rTMS causes neuroplasticity, axon growth and documented long-lasting effects in neurological patients (*86, 87*) remain to be defined, and likely involve fundamental genetic reprogramming by pioneer transcription factors.

We have begun testing our HF-rTMS protocol on human subjects. Initial results indicate that the HF-rTMS procedure is well tolerated by able-bodied volunteers and chronic SCI patients. Most interestingly, one SCI patient experienced significant and long-lasting improvement of hand function (*88*). This study is ongoing. If successful, HF-rTMS could emerge as a non-invasive, low risk treatment option to facilitate axon regeneration, alone or in combination with additional interventions, for SCI or other patients who could benefit from CNS circuitry repair.

## MATERIALS AND METHODS

### Animal models

All procedures were performed in compliance with animal protocols approved by the IACUC at Weill Cornell Medicine, New York. Histological and behavioral analyses were conducted in a blinded manner.

### Conditional gene targeting

Mouse lines *lsl-kaBaf* (*21*), *lsl-tdTom* (*20*), *lsl-EgfpL10* (*19*), *Fezf2-CreER^T2^* (*18*), *Mek1 ^f/f^* (*90*), *Mek2^-/-^* (*91*), *lsl-kaBraf* (*21*) and *Pten ^f/f^* (*92*) on a mixed 129Sv and C57BL/6 background, both males and females of matched ages, were used for this study. Mouse breeding and PCR genotyping were performed as described previously (O’Donovan et al., 2014). To induce Cre recombination in CSNs, mice aged 5 – 6 weeks and harboring the *Fezf2-CreER* allele were treated with 50 mg/kg body weight tamoxifen (Sigma-Aldrich,T5648) by oral gavage for 5 consecutive days. Surgery was performed 7 days after the last tamoxifen dose.

### Tissue collection and RNA extraction

One week after the last tamoxifen administration, Mice were euthanized by IP injection of a lethal dose of ketamine/xylazine followed by cervical dislocation. 20-25mg sensorimotor cortex tissue harboring corticospinal projection neurons was collected for TRAP-seq. RNA preparation was performed as described in Heiman et al. (2014). One mouse was used for each biological replicate.

### RNA-seq and data analysis

RNA library preparations, sequencing reactions, and initial bioinformatic analysis were conducted at GENEWIZ, LLC. (South Plainfield, NJ, USA). RNA samples were quantified using Qubit 2.0 Fluorometer (Life Technologies, Carlsbad, CA, USA) and RNA integrity was checked using Agilent TapeStation 4200 (Agilent Technologies, Palo Alto, CA, USA). RNA sequencing libraries were prepared using the NEBNext Ultra II RNA Library Prep Kit for Illumina using manufacturer’s instructions (NEB, Ipswich, MA, USA). Briefly, mRNAs were first enriched with Oligo(dT) beads, then fragmented for 15 minutes at 94 °C. First strand and second strand cDNAs were subsequently synthesized. cDNA fragments were end repaired and adenylated at 3’ends, and universal adapters were ligated to cDNA fragments, followed by index addition and library enrichment by limited-cycle PCR. The quality of the sequencing libraries was validated on the Agilent TapeStation Bioanalyzer (Agilent Technologies, Palo Alto, CA, USA), and quantified using the Qubit 2.0 Fluorometer (Invitrogen, Carlsbad, CA), as well as by quantitative PCR (KAPA Biosystems, Wilmington, MA, USA). The sequencing libraries were clustered on flowcells which were loaded onto the Illumina HiSeq instrument (4000 or equivalent) according to manufacturer’s instructions. The samples were sequenced using a 2x150bp Paired End (PE) configuration. Image analysis and base calling were conducted by the HiSeq Control Software (HCS).

Raw sequence data (.bcl files) generated from Illumina HiSeq were converted into fastq files and de-multiplexed using Illumina’s bcl2fastq 2.17 software. One mismatch was allowed for index sequence identification. Raw reads were quality checked with FastQC v0.11.7 (http://www.bioinformatics.babraham.ac.uk/projects/fastqc/), and adapters were trimmed using Trim Galore v0.6.7 (http://www.bioinformatics.babraham.ac.uk/projects/trim_galore/). The trimmed reads were mapped to the mouse reference genome (GRCm38) using STAR aligner v2.7.6a in two-pass mode. Gene abundances were calculated with featureCounts from the Subread package v.1.6.2 using composite gene models from Gencode vM25. Only unique reads that fell within exon regions were counted. The gene abundance table was filtered to retain only genes that had at least 10 reads total across all samples. The filtered abundance table was used for downstream differential expression analysis. Differentially expressed genes were determined with DESeq2 v1.32.0 using Wald tests (q < 0.05) with a two-factor model incorporating batch. Genes with an adjusted p value of below 0.05 were considered statistically significant. Principal component analysis was performed using the plotPCA function from DESeq2 v1.32.0, after removing batch effects with limma v3.48.0 removeBatchEffect. Expression heatmaps were generated using batch-corrected DESeq2-normalized log2 counts, with the values centered and scaled by row. The scripts used for principal component analysis, differential expression analysis, and heatmap generation are available at the following GitHub repository: https://github.com/abcwcm/Zhong2022.

Gene ontology analysis was performed using Ingenuity Pathway Analysis (IPA, Qiagen, Maryland) according to manufacturer’s instructions. Published datasets from sciatic nerve injury (GSE30165, GSE65053, *35*), GSE26350 (*25*), spinal cord injury (GSE126957, *47*) and kaB-RAF mediated gene expression in intestinal epithelial cells (GSE106330, *72*) were used for IPA analysis and comparison in addition to the dataset reported in this paper.

### *In situ* hybridization

Animals were administered a lethal dose of anesthetics followed by transcardial perfusion with 4% paraformaldehyde. Brains were dissected and post-fixed in the same fixative at 4°C, then incubated for 2 days in 30% sucrose solution in PBS (pH7.0) at 4°C. Samples were embedded into O.C.T. compound (Tissue-Tek) and snap-frozen on dry ice, then serially sectioned (20 μm thickness) on a cryostat (Leica CM1860). The sections were mounted on slides (Superfrost Plus 6776214 from Thermo Scientific) which were baked for 30mins at 60°C. From here on, we followed the RNAscope protocols (RNAscope™ Fluorescent Multiplex Assay) according to manufacturer’s instructions. Kits used were RNAscope Multiplex Fluorescent Reagent Kit v2 (ACD #323100), RNAscope Probe Diluent (ACD #300041), RNAscope 3-plex negative control probes (ACD #320871), and the Opal dyes Opal 650 (PerkinElmer #2553339), Opal 570 (AKOYA Biosciences #OP-001003) and Opal 520 (AKOYA Biosciences #OP-001001)

### Unilateral pyramidotomy (uPx)

The procedure was performed as described in Liu, Lu, Lee, Samara, Willenberg, Sears-Kraxberger, Tedeschi, Park, Jin, Cai, Xu, Connolly, Steward, Zheng and He (*44*). Briefly, following an incision at the side of the trachea contralateral to the dominant forelimb and exposure of the skull base, the occipital bone was cut to access the medullary pyramid. The pyramid was then cut medially with a fine scalpel (Micro Ophthalmic Scalpel, Feather) up to the basilar artery.

### Dorsal hemisection (dHx)

T8 dorsal hemisection surgery was performed as previously described (*44, 93, 94*). Briefly, following a midline incision over the thoracic vertebrae and laminectomy at T8, the dorsal spinal cord was cut to a depth of 0.8 mm with a pair of microscissors (Vannas-Tübingen Spring Scissors No. 15003-08, FST). A fine microknife (Feather micro scalpel #200300715, PFM Medical) was then drawn across the entire dorsal aspect of the spinal cord to the same depth to ensure a complete injury. The muscle layers were sutured and the skin secured with EZ clip wound closures (Stoelting), or absorbable 4-0 sutures (Ethicon) for mice later treated with HF-rTMS.

To check for the completeness of dHx lesions, spinal transverse sections 5 mm caudal and rostral from the lesion center were prepared and stained for BDA or PKCγ (Abcam, ab109539). For mice with uPx, we collected transverse sections at level C1 and stained them for PKCγ.

### Anterograde labeling of CST axons

Two weeks before terminal perfusion, biotinylated dextran amines (BDA 10%, Thermo Fisher Scientific, 10,000 MW, Lysine Fixable) were administered as described by Zukor, Belin, Wang, Keelan, Wang and He (*95*), with modifications. Briefly, fine glass pipets were pulled with a micropipette puller (Sutter Instruments P-97) from borosilicate glass capillaries (WPI). The mouse head was stabilized in a stereotaxic frame (Kopf, Model 900), and a unilateral 1 mm x 3 mm window was cut from the skull over the sensorimotor cortex using a microdrill (Foredom, Portable Rotary Micromotor Kit). 400 nl of BDA were injected at a rate of 80 nl/min and a depth of 0.7 mm into each of 6 injection sites with a microinjector (WPI, Nanoliter 2010) using the fine glass pipets. For mice subjected to dHx we injected at anteroposterior (AP) coordinates from bregma (in mm): -0.5/1.2, - 1.0/1.2, 0/1.5, −1.5/1.2, −0.5/1.8 -1.0/1.8, -1.5/1.8. For bilateral tracing, a similar cranial window was cut at the contralateral side, sparing the midline. For uPx we used the following coordinates for BDA injection (mm): 1.8/0.8, 1.8/1.3, -0.2/1.5, −0.7/1.5, −0.2/2.0, -0.7/2.0, depth 0.7 mm from the pial surface (*96*). The skin was closed with either metal wound clips, or absorbable 4-0 sutures for mice subjected to HF-rTMS. To detect BDA labeled axons, tissue sections were incubated with 0.6% H_2_O_2_ in PBS for 15 minutes, then incubated in streptavidin-HRP (Perkin Elmer, NEL750001EA) in PBS for three hours at room temperature. From there on, procedures were performed following manufacturer’s instructions (TSA kit, Perkin Elmer).

### Anterograde transsynaptic tracing of corticospinal connectivity

AAV2/1-WGACreER^T2^ (2.5 × 10^13^ GC/ml, UPenn Vector Core) was unilaterally injected into cortex of *lsl-tdTom+* mice 14 days prior euthanasia as described above for BDA. Tamoxifen treatment began 5 days later. For quantitation of synaptic connection, sections were stained for tdTom; tdTom positive cells were counted on three adjacent sections. Efficiency of transsynaptic tracing was tested in non-injured mice 10 days after AAV injection (**Suppl. Figs. 6A,B**). For whole-mount visualization of labeled interneurons, we adapted the iDISCO+ protocol (*97*), **Suppl. video clip 2**). Tissues were delipidated with a methanol gradient (25%, 50%, 75%, 100%), followed by reverse gradient extraction, then washed with 0.1% Triton X-100/0.3 M glycine for 1h twice, followed by PBS/0.1% Triton X-100/0.05% Tween 20/2 μg/ml heparin (PTxwH buffer) for 1hr twice. Samples were then incubated for 3 d with anti-RFP primary antibody (Rockland, #600-401-379), and subsequently for 3 d with secondary Ab (anti-rabbit secondary from Molecular Probe or Jackson Immunology) in PTxwH, followed by dehydration in a methanol gradient (25%, 50%, 75%, 100%), then two 30 min incubations in dichloromethane (100%) and finally an overnight clearing step in dibenzyl ether (DBE; Sigma-Aldrich). 16 bit image files were acquired on the light sheet microscope with an sCMOS camera (LaVision Biotec Ultramicroscope II) and processed using ImageJ for level adjustment and maximum projection, Arivis (Vision4D, Arivis) for alignment and stitching, and Imaris (Bitplane) for 3D rendering, to visualize and quantify spinal interneuron distribution.

### Ablation of corticospinal neurons after regeneration

400 nl HiRet lenti virus (2 × 10^12^ GC/ml, HiRet-FLEX-DTR for ablation, HiRet-GFP for control, Boston Children’s Hospital Viral Core) were injected into the spinal cord bilaterally (0.35 mm lateral, 0.5 mm depth) 2 mm caudal to the epicenter of T8 dHx 8 weeks after lesion. Animals were tested for horizontal ladder walking to assess their skilled stepping performance 2 weeks post injection. Diphtheria toxin (DT, Sigma D0564, 100 ug/kg) was then administered i.p. to all animals, and they were re-tested on the horizontal ladder 2 weeks later.

### Retrograde labeling or targeting of T8 projecting CSNs

Mice were bilaterally injected in the spinal cord gray matter at T8, 3 injections (500 µm apart) each side and 400 nl AAV per injection. Biotin-conjugated Cholera Toxin B subunit (CTB, Sigma-Aldrich, C9972) or AAVrg-CAG-GFP (Addgene viral prep 37825-AAVrg) was used for retrograde labeling of T8 projecting neurons and AAVrg-pgk-Cre (Addgene viral prep 24593-AAVrg) was used to retrogradely targeting T8 projecting neurons. AAV titers were ≥ 7×10¹² vg/ml. AAVrg-CAG-GFP was a gift from Edward Boyden. AAVrg-pgk-Cre was a gift from Patrick Aebischer.

### High Frequency rTMS

HF-rTMS was delivered using a MagPro X100 (MagVenture) stimulator connected to a 40 mm diameter single circular coil (MagVenture, Cool-40 Rat). Coil temperature was maintained at 37°C. Mice received one HF-rTMS session daily, starting two days after the SCI surgery, as follows: The awake mouse was restrained in a transparent DecapiCone (Braintree Scientific, DC M200) and position in a mouse holder (Colonial Medical Supply, MR01C). The coil was placed over the skull, with the coil center positioned over Lambda. The focal plane of the coil was parallel to the calvarial bone and the longitudinal axis aligned with the midline. The default settings of stimulator were diphasic (full-sine) waveform with a 280 microseconds pulse width. Resting motor threshold (RMT), the minimum stimulus intensity that evoked a twitch of the hindlimb (*98*), (*99*), was determined for each mouse before the first treatment session by monitoring the muscle twitch in response to single TMS pulses. The stimulator output evoking RMT was used as reference for all subsequent treatment sessions; this was typically around 35% of maximum stimulator output (MSO, maximum initial dB/dt = 80 kT/s). Each experimental animal received one daily HF-rTMS session comprising 750 pulses (75 pulses per train, 10 repetitions with 60 seconds intervals, at a frequency of 15 Hz and an intensity of 120% RMT. The mouse was awake during the entire procedure and could breathe normally through an opening in the tip of the plastic cone. Sham stimulation was applied using the same parameters, but with the coil placed perpendicular to the skull to prevent magnetic stimulation of the brain. A piece of Mu-metal sheet was positioned between the tilted coil and the skull to assure delivery of the coil’s vibration to the mice, while no TMS-induced muscle twitch was elicited. Mice were placed back to the cage after the treatment.

### Small animal TMS model

We utilized the Digimouse dataset (*100*) to approximate the volume conductor profile of the adult mice. The pre-existing segmentation of the imaging data was utilized to extract the boundary surfaces separating skin, skull, and brain as this has been shown to result in a reasonable approximation in the case of human TMS modeling (*101*). The dimensions of the Digimouse dataset were uniformly scaled to match the size of the animals used in the study. The geometric wire winding pattern of the small animal TMS coil (MagVenture Cool-40 Rat) was modeled based on specifications obtained from the from the manufacturer (MagVenture). The coil model was validated by comparing the predicted values of *∂**B***/*∂t* against reference measurements provided the coil manufacturer. For simulating the induced E-field distributions, the coil was positioned on the Digimouse dataset to the corresponding location used for the empirical stimulation, and the TMS coil current amplitude values were matched to the empirical stimulator output intensity values (rMT) utilizing the known linear relationship between these two quantities.

### Computing the TMS-induced E-fields

The TMS-induced E-field is determined by the Maxwell–Faraday law of induction ∇ × ***E*** = −*∂**B***/*∂t* from which it follows that the total E-field has the general form ***E*** = −*∂**A***/*∂**t*** − ***∇***ϕ. The first term involving the vector potential ***E***^***inc***^ = −*∂**A***/*∂t* corresponds to the primary field (excitation) induced by the current in the coil, whereas the second term involving the scalar potential ***E***^***S***^ = −***∇***ϕ ( the secondary field) arises from the differences in the electrical conductivities of the tissue compartments (see, *e.g.,* (*102*). The primary and secondary fields are coupled through the quasi-static condition ∇ · ***J*** = ∇ · (*σ**E***) = 0. For the numerical solution of the TMS-induced E-fields we adhere to our recently proposed boundary-element method (BEM) that operates directly with electric charges associated with ϕ and can be made computationally highly efficient using the fast multipole method [FMM, (*62*)]. Briefly, we discretize the total combined surface *S* of all conductivity boundaries (tissue classes) into *N* small planar triangles *t*_*i*_, *i* = 1, …, *N* with centers ***c***_*i*_, *i* = 1, …, *N*, outer normal vectors of every manifold tissue shell ***n***_*i*_, *i* = 1, …, *N*, and triangle areas ***A***_*i*_, *i* = 1, …, *N*. The unknown surface charge density is constant for every triangle and equal to *ρ*_*i*_, *i* = 1, …, *N*. Purely parametric time dependence due to the quasi-static conditions is eliminated via separation of variables. This formulation results in a system of linear equations for unknown induced charges at the boundaries solved iteratively. For the *n*:th iteration,

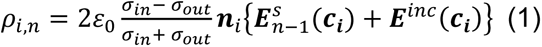

and

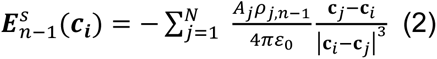

where ***E***^*inc*^ is the excitation field (created by a coil), ***E***^***s***^ is the secondary field of charges induced at the interface of any two compartments with conductivities *σ*_*in*_ and *σ*_*out*_, respectively, and *ε*_0_ is permittivity of vacuum (1). The major burden of the iterative solution is the repeated computation of the electric field from a large ensemble of point charge sources at a large number of target points following Eq. (2) which is highly effectively handled by the FMM algorithm.

### Horizontal ladder-walking test

For all tasks, mice were handled and exposed to the behavioral assay settings for one week prior to the surgery, to collect baseline data and to eliminate mice with any obvious behavioral abnormalities. Testers were blinded with respect to treatment groups. The customized horizontal ladder apparatus consists of two transparent Plexiglas walls of 1 meter in length and irregularly placed removable metal rungs. The smallest distance between two rungs was 1 cm. Prior to surgery mice were habituated to the apparatus and trained twice daily, every two days, for 6 days. Baseline values for all mice were collected the day before surgery. Mice with persistent high baseline error rate ( > 30% than the normal rate) were excluded from the experiments. After surgery, animals were tested weekly for the first two weeks and biweekly thereafter. Each mouse walked on the irregular horizontal ladder forward and back for a total of 4 times. The number of hind limb misplacements was counted. A completely missed step, slipping, or mis-placement followed by re-placement of the paw were counted as errors. The number of errors per trial was presented as percentage of error compared to the total number of steps taken during the trial.

### Pellet grasping test

Mice were food-restricted to maintain 80% -85% of their *ad libitum* feeding weight. For habituation and training, and to determine each mouse’s dominant paw, all mice underwent three training sessions prior to surgery, retrieving sugar pellets presented within reaching distance outside a transparent Plexiglas chamber (Supplementary video 2). uPx was then performed on the contralateral side to denervate CST projecting to the dominant paw. Mice were tested 1, 3 and 6 weeks after uPx. The slip of the Plexiglas chamber was positioned the way that only the dominant paw could be used to grasp the food pellet. Forepaw movements were videotaped and the retrieval success rates quantified.

### Patch removal test

Precut tape (6 mm x 6 mm, 3M, Office Depot) was applied to one forepaw, alternating between affected and unaffected ones. The time a mouse took to remove the tape was recorded. For habituation and training prior to surgery, mice were subjected to this procedure twice daily on every second day of 6 days, for a total of 6 sessions. Mice persistently taking longer than 2 min to remove the tape were excluded from the experiments. Four days, 3 and 6 weeks post surgery mice were tested for tape removal as follows: tape was applied to the forepaw, alternating between right and left, three times each, with 20 minute intervals between tests. Mice were videotaped until they had removed the tape, or for a maximum of 3 min. The time elapsing from the mouse first noticing the tape to its removal was recorded, and averaged for each side.

### Western blot analysis

Cortices of HF-rTMS treated or sham treated mice were dissected within 20 min of the last HF-rTMS treatment, and immediately frozen in liquid nitrogen. Protein extractions were performed as described previously (*7*). Protease inhibitor and phosphatase inhibitor cocktails (Sigma-Aldrich, P8340 and P0044) were added to the lysis buffer to prevent protein degradation and dephosphorylation. An equal amount of total protein (30 μg) was loaded in each lane. Primary antibodies were anti-MEK1/2 (Cell Signaling, 4694), anti-pMEK1/2 (Cell Signaling, 9121), anti-AKT (Cell Signaling, 2920), anti-pAKT (Cell Signaling, 9271) and anti-βIII tubulin (Cell Signaling, 5666) antibodies. Signals were detected and quantified using an Odyssey Infrared Imaging System (LI-COR). Protein levels were normalized to that of βIII-tubulin. Protein levels of sham-treated mice were set to an arbitrary unit of one and protein levels in kaB-RAF animals are relative to that within each wild type tissue.

### Histology and immunohistochemistry

Animals were administered a lethal dose of anesthesia followed by transcardial perfusion with 4% paraformaldehyde. Brains and spinal cords were dissected and post-fixed in the same fixative at 4°C, then incubated overnight in 30% sucrose solution in PBS (pH7.0) for cryoprotection. Samples were embedded into O.C.T. compound (Tissue-Tek) and snap-frozen on dry ice, then sectioned (25 μm thickness) using a cryostat (Leica CM1860). To evaluate regeneration after dHx serial sagittal sections spanning 5 mm proximal through 5 mm distal of the lesion site were collected. To assess the extent of CST sprouting after uPx, three C5 serial transverse sections were analyzed.

Immunostaining was performed following standard protocols as described previously (*10*). All antibodies were diluted in PBS containing 10% normal goat serum (NGS, ThermoFisher Scientific) and 0.3% Triton X-100 (Sigma-Aldrich). Sections were incubated with primary antibodies overnight at 4°C. Primary antibodies used were rabbit anti-GFAP (Millipore AB5804), or goat anti-RFP (Rockland, 600-401-379). PKCγ (Abcam, ab109539). Alexa Fluor conjugated secondary antibodies (donkey-anti-rabbit Alexa-488, ThermoFisher Scientific, C22841, donkey-anti-rabbit Alexa-555, Thermo Fisher Scientific, A31572, donkey-anti-goat Alexa-488, Abcam ab150133, and rabbit-anti-goat Alexa-594, Abcam, ab150148) were then applied at a dilution of 1:500 and incubated for 3 h at room temperature.

### Confocal imaging

Images were acquired using a Zeiss LSM-710NLO microscope and processed using the manufacturer’s ZEN software.

### Quantification of axon regeneration

For mice subjected to dHx, numbers and lengths of regenerating axons were counted and measured as described previously (*44, 103*) . In brief, transverse sections of the medullary pyramid of medulla pyramid 1 mm proximal to the pyramidal decussation were used to count BDA-labeled CST fibers for normalization. A series of rectangular segments 100 μm wide as high as the dorsal-ventral aspect of the cord were superimposed onto the sagittal sections, starting from 1 mm rostral up to the lesion center. The number of intersections of BDA-labeled fibers with a dorsal-ventral line positioned at a defined distance caudal from the lesion center was counted. Fibers were counted on 3 sections with the main dorsal CST. The number of counted fibers was normalized by the number of total BDA-labeled CST fibers in the medulla and divided by the number of evaluated sections. This resulted in the number of CST fibers per labeled CST axons per section at different distances (fiber number index).

### Quantification of axon sprouting

For mice subjected to uPx, axon sprouting was quantified from C7 spinal cord transverse sections by densitometry and sprouting length similarly to (*44*). For densitometry measurements, the pixel densities of the two sides of the gray matter were measured using ImageJ, sub-thresholded to the background and normalized by area. The sprouting density index was then calculated as the ratio of contralateral to ipsilateral counts. Three sections were measured for each mouse. To quantify the length of sprouting axons, horizontal lines were drawn on images through the central canal and across the lateral rim of the gray matter. Three vertical lines (Mid, Z1, and Z2) were then drawn to divide the horizontal line into three equal parts from the central canal to the lateral rim. While Mid denotes midline crossing fibers, Z1 and Z2 indicate sprouting fibers at different distances from the midline. Fibers crossing each of the three lines were counted on each section and normalized to the total number of CST fibers as counted at the medulla level. Three sections were counted for each mouse.

### Quantification of post-uPx synaptogenesis

The total number of tdTom^+^ cells was counted for each animal at the cervical level 5-6. At least three sagittal sections from each animal were counted. The total number of tdTom+ cells was quantified on the ipsilateral and contralateral sides and divided by the total number of sections analyzed.

### Statistical analysis

GraphPad Prism (GraphPad Software) was used for statistical analyses. Data were analyzed using ANOVA and the Bonferroni within-groups comparison with additional testing using Dunnett’s test or Student’s t test. Values of P < 0.05 were considered significant. All error bars indicate SEM. Two-tailed Student’s t-test was used for the single comparison between two groups. Other data were analyzed using one-way or two-way ANOVA depending on the experimental design (see figure captions). *Post hoc* comparisons were carried out only when a main effect showed statistical significance. P-values of multiple comparisons were adjusted by using Bonferroni’s correction. Data are presented as means ± SEM; asterisks indicate statistical significance under an appropriate test.

## Supplementary Materials

**Figure S1:**
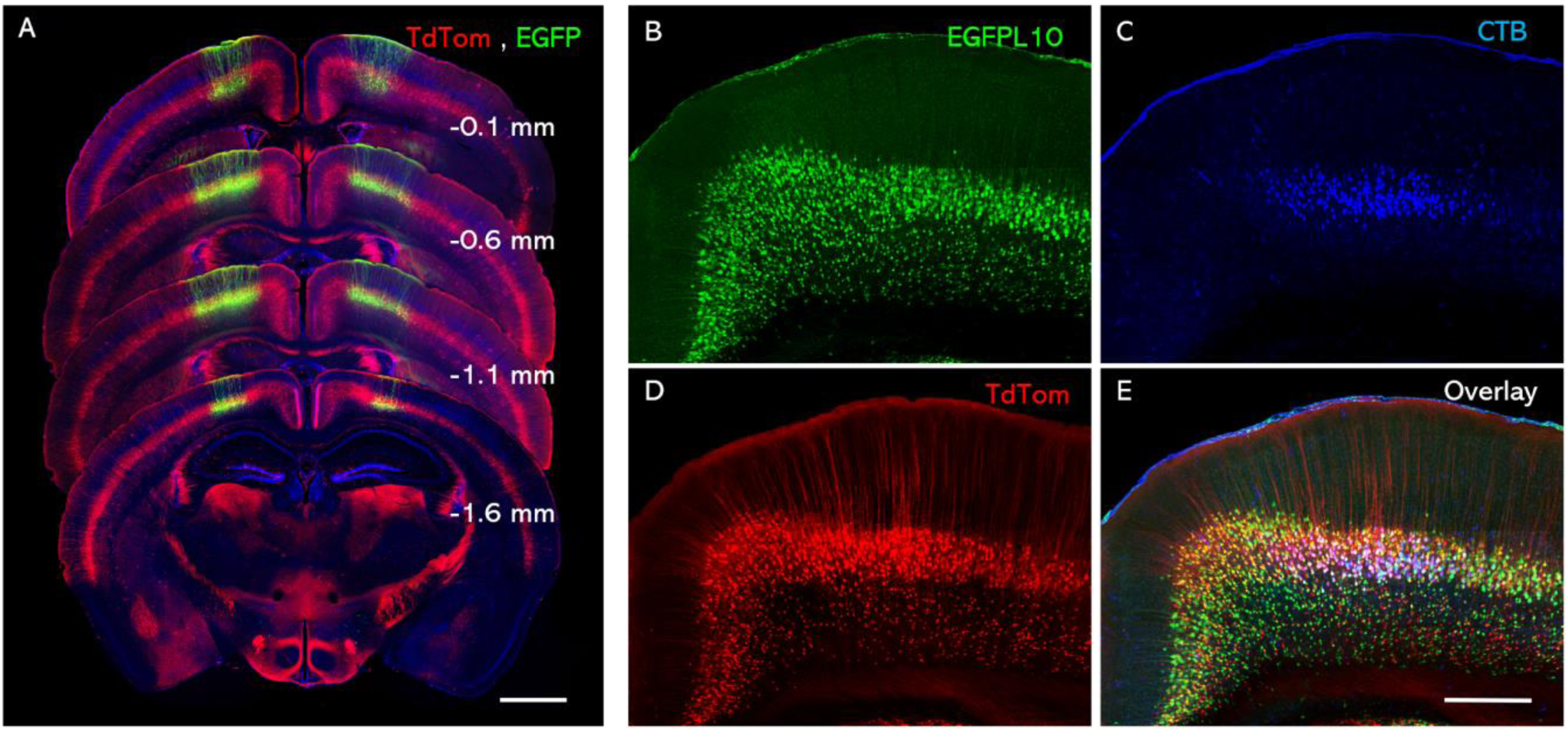
Retrograde labeling identified FEZF2+ cells as corticospinal projection neurons. (A) AAVrg-EGFP was injected into the spinal cord of a tamoxifen treated *lsl-tdTom :* a *Fezf2-CreER* mouse at T8 bilaterally, to retrogradely label T8-projecting CSNs. The numbers indicate the distance of each section from bregma. Red, tdTom; Green, EGFP. Blue, Draq5. Scale bar, 100 µm. (B-E) Cholera Toxin B (CTB) injected into spinal cord at T8 retrogradely labeled FEZF2+ neurons in sensorimotor cortex. In *lsl-EgfpL10 : lsl-tdTom : Fezf2-CreER* mouse treated with tamoxifen, 93.0 ± 4.9 % CTB positive cells concomitantly express EGFP and or tdTom (mean ± SEM, N=3.)

**Figure S2.**
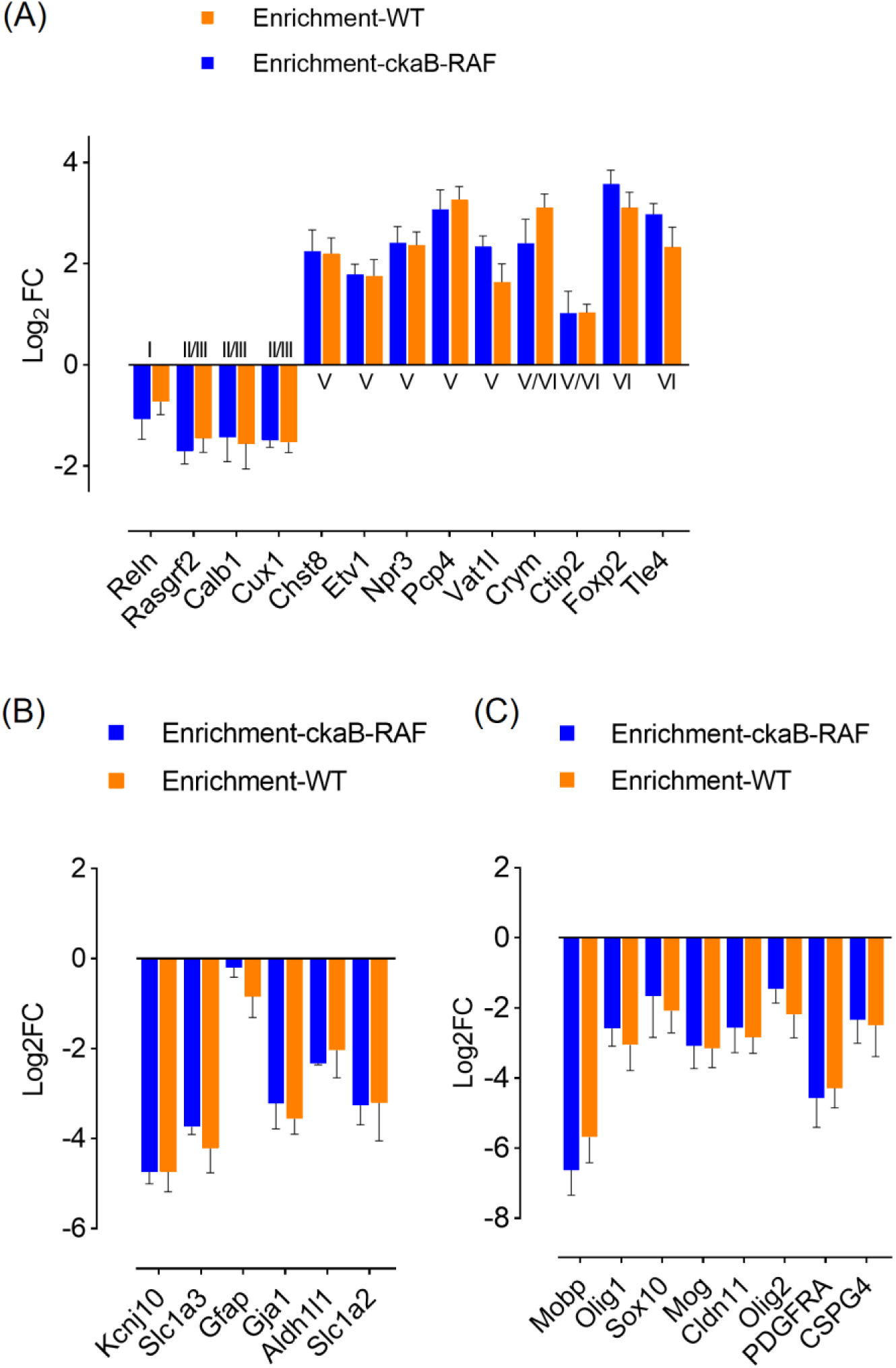
Enrichment of transcripts by TRAP from mice harboring the *Egfpl10 : Fezf2-CreER* allele. (A) Enrichment of layer V and VI specific neuronal markers. (B) Decrease of astrocyte and (C) oligodendrocyte markers. Mean ± SD. N = 3 biological replicates of RNA-seq or TRAP-seq per condition. p < 0.005, Wilcoxon matched-pairs signed rank test.

**Figure S3:**
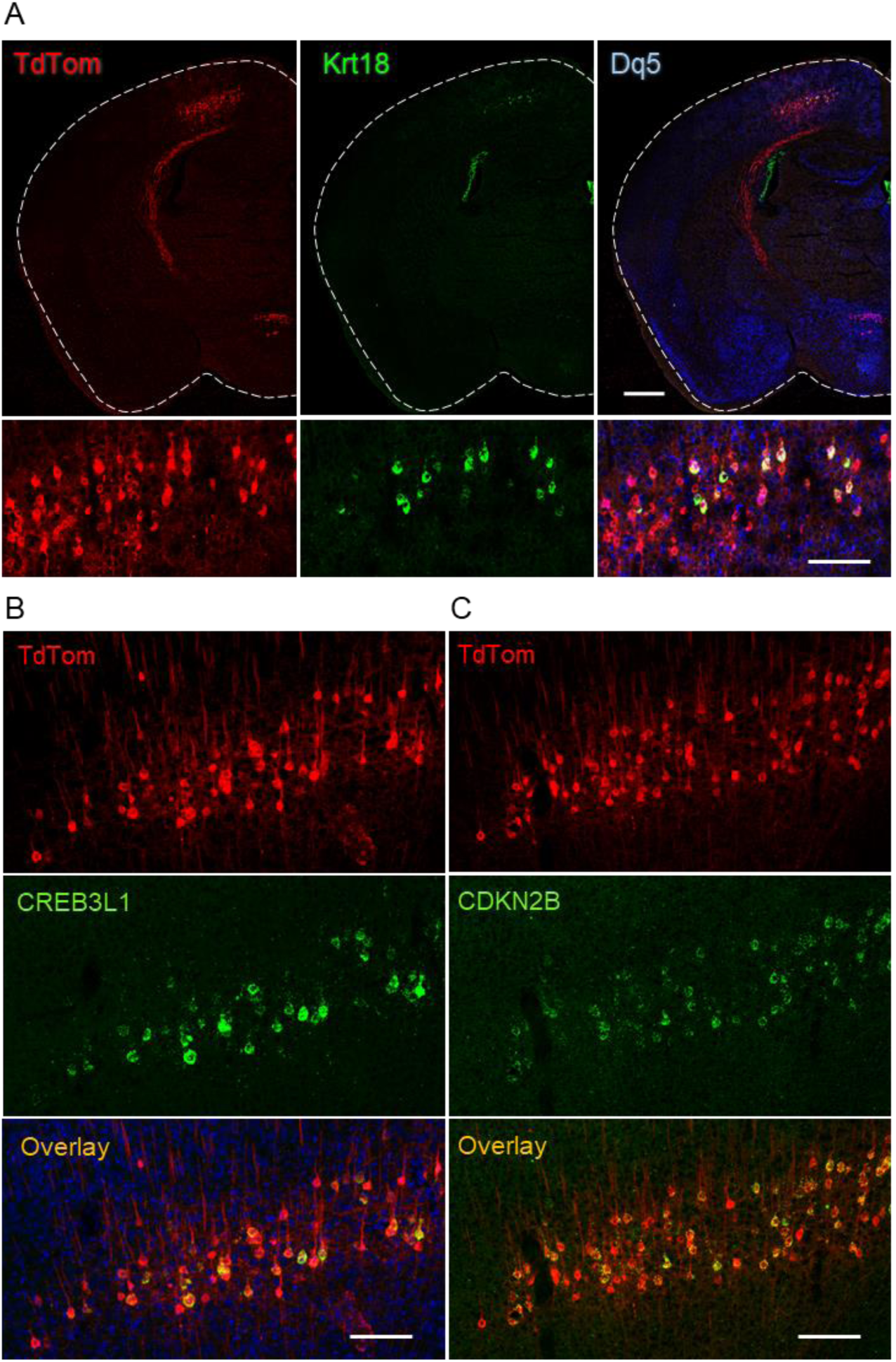
Verification of B-RAF induced transcripts by *in situ* hybridization (green). AAVrg-Cre was injected into T8 spinal cord of *an lsl-kaBraf:tdTom* mouse to induce kaB-RAF and tdTom (red) expression in CSNs. The animal was perfused 7 days later and sections stained with specific antisense probes and anti-tdTom. (A) intermediate filament keratin-18, (B) transcription factor CREB3L1, (C) cyclin-dependent kinase inhibitor CDKN2B. Scale bars, (A) top, 1mm, bottom, 250 µm. (B, C) 250µm.

**Figure S4:**
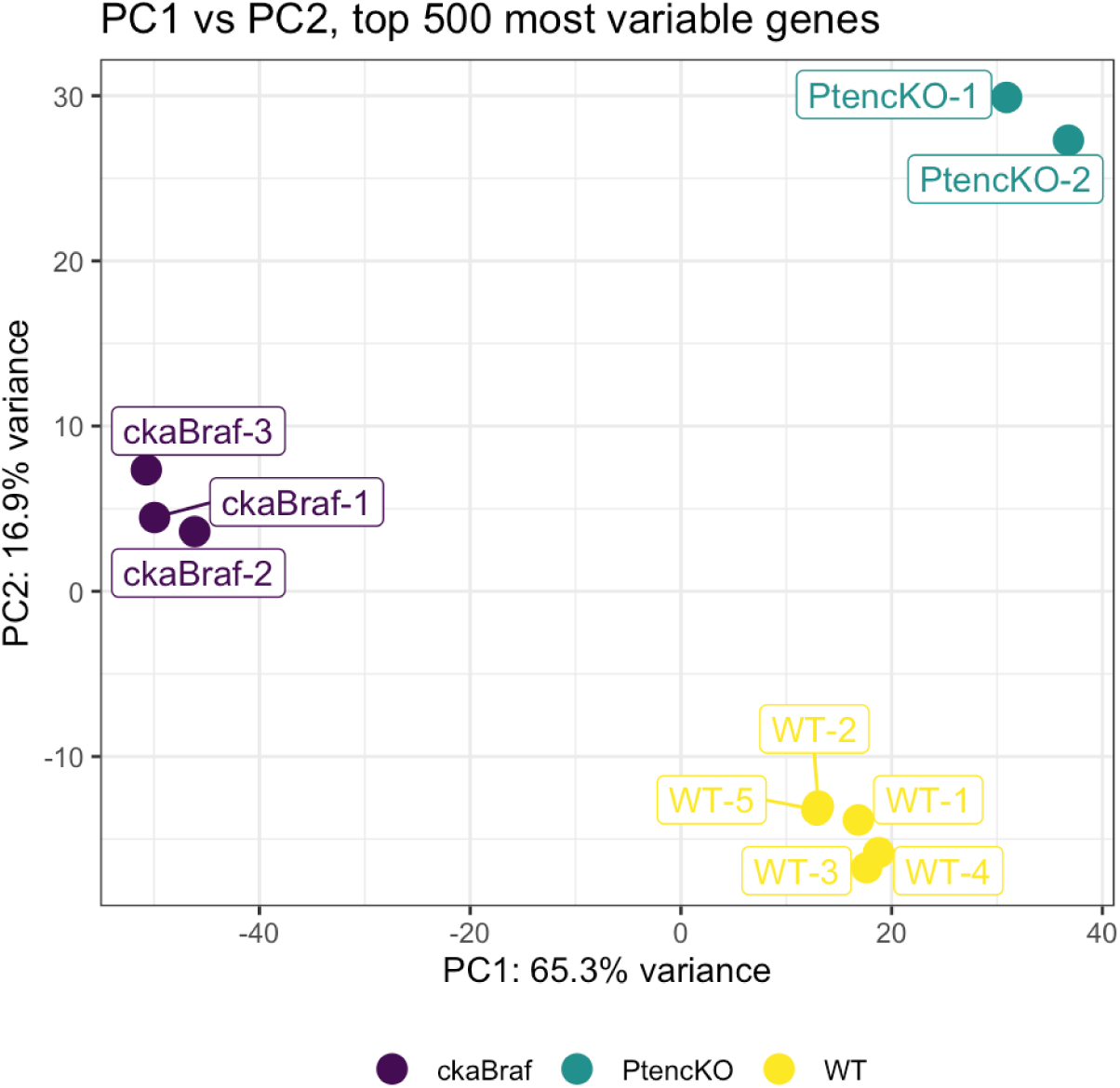
Principal component analysis (PCA) comparing TRAP-seq DEGs induced in CSNs by kaB-RAF expression (N = 3, green) or by PTEN deletion (N= 2, purple) with baseline gene expression in CSNs (N = 5, yellow). PCA was performed using the top 500 most variable genes, which were computed on batch-corrected DESeq2-normalized log2 counts. Batch effects were corrected with the remove Batch Effect function from the limma package v3.48.0.

**Figure S5:**
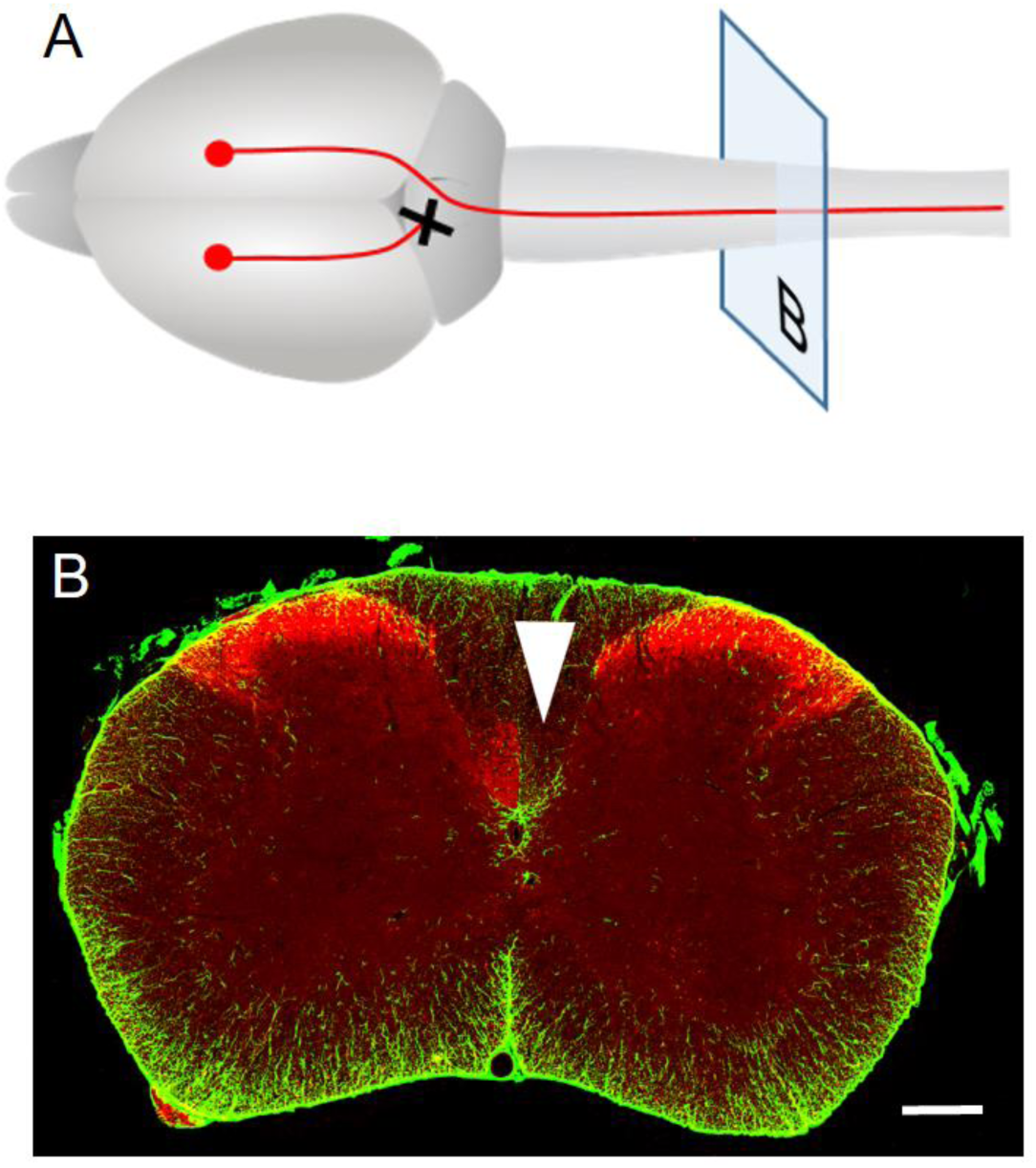
Unilateral pyramidotomy completely interrupts CST axonal connection. (A) Illustration of unilateral pyramidotomy paradigm. (B) Representative transverse section of cervical spinal cord showing absence of PKCγ immunoreactivity in the transected CST (white arrow). Tissues were analyzed 6 weeks after pyramidotomy. Red, PKCγ; Green, GFAP. Scale bar, 150 μm.

**Figure S6:**
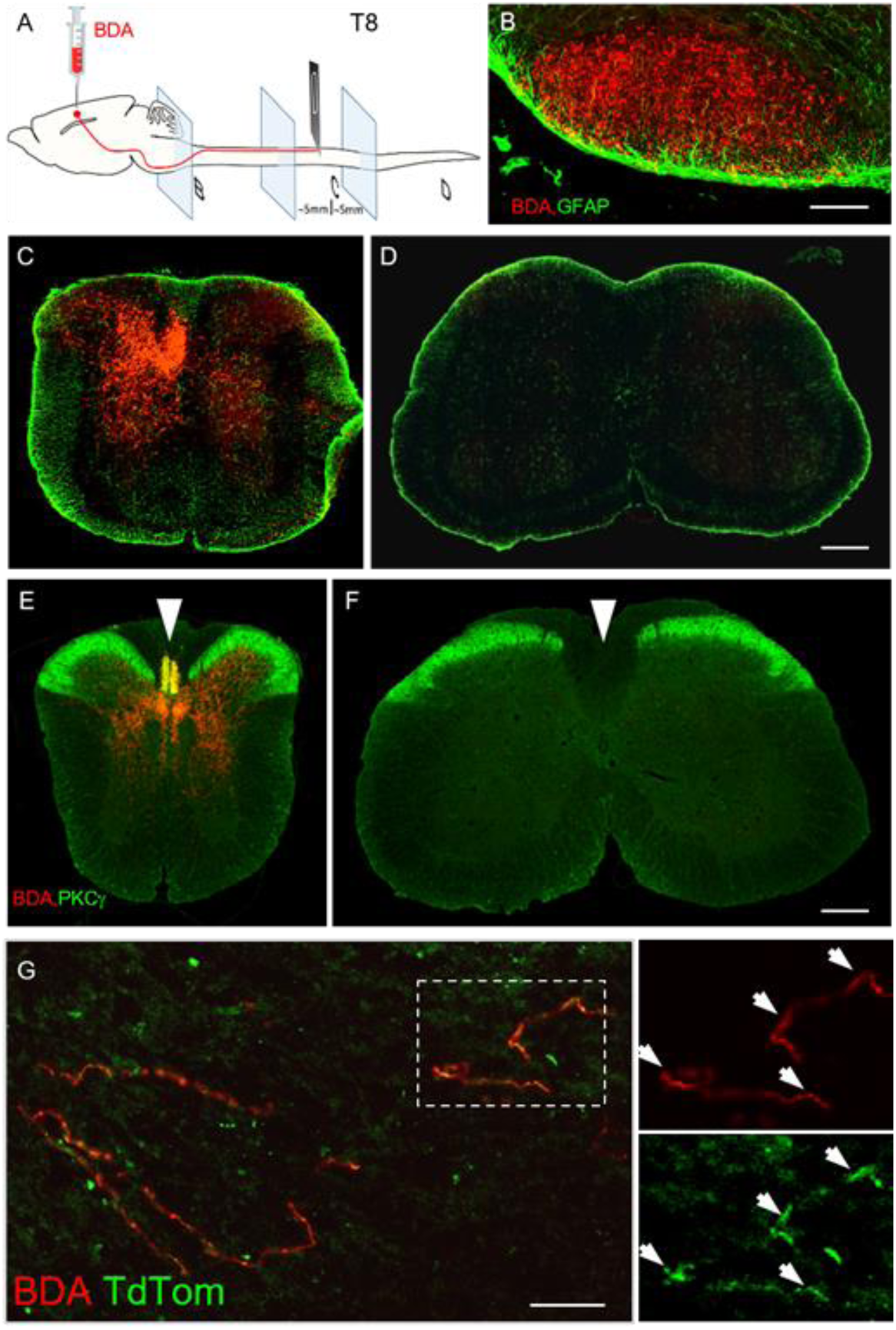
Spinal cord T8 dorsal hemisection completely interrupts CST axonal connections. (A) Schematic of experimental procedures. dHx was carried out in 7 weeks old mice. Two weeks before perfusion, BDA was injected unilaterally into sensorimotor cortex. Transverse sections show BDA staining at (B) medulla oblongata, (C) thoracic segment ∼ 5 mm rostral to the lesion epicenter, and (D) lumbar segment ∼ 5 mm caudal to the lesion epicenter. Red, BDA; Green, GFAP. (E, F) T8 dHx resulted in a complete loss of PKCγ, a marker of CST axons. BDA and PKCγ staining of transverse sections (E) rostral and (F) caudal to the lesion epicenter. White arrowheads indicate complete loss of PKC γ staining in the CST. Red, tdTom; green, PKC γ. Scale bar, 200 µm.

**Figure S7:**
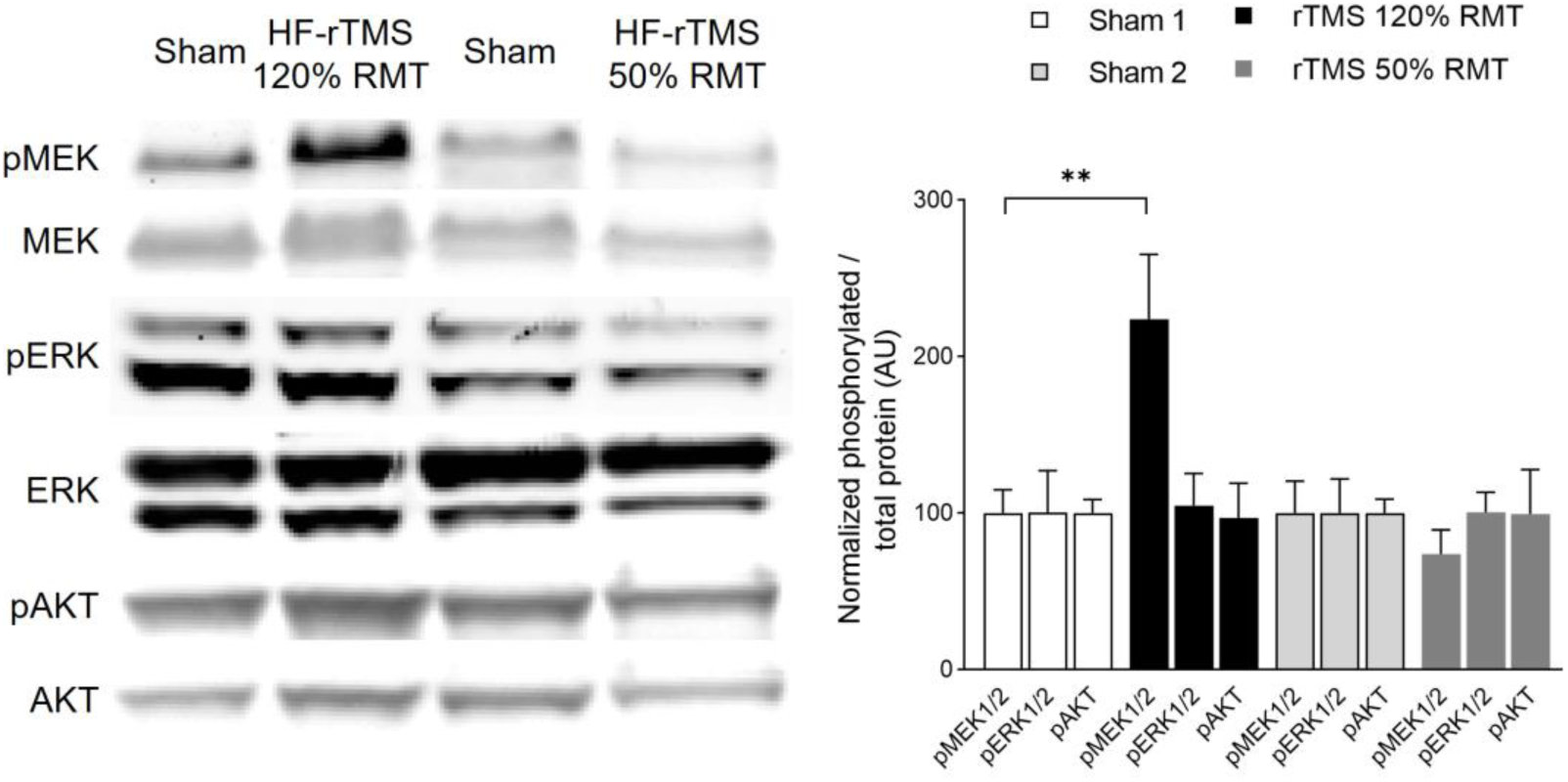
Supra-threshold HF-rTMS activates MEK1/2 kinases in mouse motor cortex. Mice were treated with 15 Hz rTMS for 5 consecutive days. Sensorimotor cortex tissue retrogradely labeled with tdTom was dissected for Western blotting. HF-rTMS at 120% of RMT increased pMEK by 120% of the baseline. No stimulation of pMEK was seen with sub-threshold HF-rTMS at 50% of RMT. AU, arbitrary unit. ** *p* < 0.001. ANOVA with Bonferroni correction. N = 3 per group. Error bars, mean ± SEM.

**Table S1:** TFs upregulated by B-RAF activation were predicted as top regulators of the gene expression program. In silicon EnrichR and ChEA3 analysis used 1091 transcripts upregulated by B-RAF activation (p_adj_ < 0.05, 1147 upregulated genes minus 56 TF transcripts) were used for the analysis. Sixty one percent (34 of 56) TFs were predicted by EnrichR with p_adj_ <0.05 and ranked as top 200 TFs predicted by ChEA3.

| Gene symbol | log<sub>2</sub> FC | *padj* | EnrichR *padj* | ChEA3 score | TF family |
| --- | --- | --- | --- | --- | --- |
| Aff1 | 0.80 | 1.12E-04 | 5.76E-05 |  | AF-4 |
| Aff3 | 0.55 | 4.75E-05 | 2.59E-05 |  | AF-4 |
| Atf6 | 0.37 | 3.35E-03 | 7.97E-07 | 493.3 | TF_bZIP |
| Bhlhe22 | 0.38 | 1.36E-02 |  | 277.3 | bHLH |
| Cbfb | 0.66 | 1.89E-03 | 2.59E-04 |  | CBF |
| Cebpd | 1.09 | 1.05E-08 |  | 200.6 | TF_bZIP |
| Creb3l1 | 2.86 | 6.18E-65 | 1.98E-13 | 326.8 | TF_bZIP |
| Csdc2 | 0.53 | 9.39E-05 | 2.05E-09 |  | CSD |
| Csrnp1 | 0.79 | 1.19E-05 |  | 53.5 | CSRNP_N |
| D630045J12Rik | 0.38 | 1.48E-03 |  |  | Others |
| Dmtf1 | 0.44 | 1.25E-02 | 1.12E-05 | 915 | MYB |
| E2f3 | 0.36 | 2.85E-03 | 4.75E-06 | 692.7 | E2F |
| Egr2 | 2.45 | 1.65E-04 | 2.92E-20 | 66.67 | zf-C2H2 |
| Elk3 | 0.96 | 5.64E-09 | 6.27E-18 | 63 | ETS |
| Etv4 | 1.12 | 2.04E-04 | 2.24E-17 | 381.3 | ETS |
| Fev | 3.15 | 2.23E-03 | 5.58E-02 | 547.3 | ETS |
| Fosl1 | 4.55 | 1.63E-03 | 2.92E-20 | 17.8 | TF_bZIP |
| Gatad1 | 0.33 | 2.22E-03 | 1.05E-03 |  | zf-GATA |
| Gm15446 | 0.36 | 3.75E-02 |  |  | zf-C2H2 |
| Gmeb1 | 0.39 | 1.53E-02 | 1.26E-02 | 921 | SAND |
| Grhl3 | 0.89 | 1.66E-02 | 2.29E-12 | 91.33 | CP2 |
| Helt | 3.57 | 4.32E-32 | 1.83E-01 | 1123 | bHLH |
| Hes1 | 0.65 | 1.29E-02 | 2.24E-17 | 396 | bHLH |
| Hmga1 | 0.94 | 2.91E-10 | 1.23E-04 | 763 | HMGA |
| Hmga2 | 5.96 | 1.80E-34 | 7.49E-12 | 287.3 | HMGA |
| Hmx3 | 6.17 | 3.06E-03 | 9.31E-01 | 1043 | Homeobox |
| Irf6 | 2.84 | 2.95E-24 | 5.88E-09 | 97.33 | IRF |
| Irf8 | 2.53 | 3.56E-31 | 1.22E-07 | 232.8 | IRF |
| Jun | 0.45 | 3.32E-03 | 4.53E-15 | 98.33 | TF_bZIP |
| Maff | 0.92 | 1.37E-02 | 7.91E-21 | 337.8 | TF_bZIP |
| Mitf | 1.06 | 5.30E-14 | 1.92E-06 | 271.2 | bHLH |
| Nfe2l3 | 1.78 | 2.27E-09 | 1.98E-13 | 370 | TF_bZIP |
| Phox2a | 2.92 | 7.52E-03 | 7.06E-03 | 487 | Homeobox |
| Phtf1 | 0.68 | 6.94E-05 |  |  | Others |
| Pias3 | 0.37 | 2.28E-02 |  |  | zf-MIZ |
| Pitx3 | 4.37 | 1.60E-07 | 1.83E-01 | 692.7 | Homeobox |
| Pknox1 | 0.38 | 1.19E-02 | 7.97E-07 | 663.3 | Homeobox |
| Pou3f3 | 0.54 | 1.38E-09 | 2.06E-03 | 699.3 | Pou |
| Ppard | 0.46 | 6.36E-04 | 2.29E-12 | 62.5 | THR-like |
| Prdm12 | 3.20 | 2.80E-02 | 7.06E-03 | 726.7 | zf-C2H2 |
| Rbpj | 0.32 | 4.55E-02 | 7.97E-07 | 540 | CSL |
| Smad7 | 0.66 | 3.22E-04 | 1.56E-18 |  | MH1 |
| Stat3 | 0.31 | 1.37E-02 | 5.88E-09 | 198.8 | STAT |
| Stat4 | 1.51 | 1.63E-02 | 2.59E-05 | 235 | STAT |
| Stat6 | 0.55 | 1.41E-04 | 6.73E-13 | 300.8 | STAT |
| Tead4 | 3.86 | 8.32E-24 | 3.32E-16 | 233.5 | TEA |
| Tgif1 | 1.35 | 5.80E-03 | 1.22E-24 | 527.3 | Homeobox |
| Twist1 | 2.31 | 2.12E-03 | 2.05E-09 | 123 | bHLH |
| Vezf1 | 0.25 | 4.89E-02 | 7.06E-03 | 740 | zf-C2H2 |
| Wt1 | 3.07 | 1.15E-03 | 4.67E-08 | 393.8 | zf-C2H2 |
| Ybx1 | 0.30 | 3.86E-02 | 5.28E-01 | 1385 | CSD |
| Zfp532 | 0.33 | 9.23E-03 |  |  | zf-C2H2 |
| Zfp672 | 0.27 | 4.68E-02 |  |  | zf-C2H2 |
| Zfp941 | 0.47 | 4.97E-02 |  |  | zf-C2H2 |
| Zmat3 | 0.30 | 2.37E-02 | 1.22E-07 |  | zf-C2H2 |
| Zmiz1 | 0.44 | 2.21E-05 |  |  | zf-MIZ |

**Table S2:** Gene ontology (GO) analysis of kaB-RAF-induced gene expression, and comparison with published studies. Ingenuity Pathway Analysis (IPA, Qiagen) was performed using TRAP-seq data from CSNs expressing kaB-RAF and compared with data from DRG neurons one- or 4-days post sciatic nerve injury (rat DRG, SNI) (GSE30165, *35*); facial motor neurons 1- or 4-days after facial nerve injury (FMN, FNI) (GSE162977, *28*) and CSNs growing into an NPC graft 3- and 7-days post-SCI (CSN, SCI+NPC) (GSE126957, *47*). The cutoff of transcripts used for IPA was log_2_FC > 1, *p*_adj_ (or *p*) < 0.05. Z-scores shown illustrate pathways with a *p* ≤ 0.05 and an activation z-score > 2 (*104*). Empty cells denote non-significant change in biological activity (p > 0.05 or Z < 2). Red, pathways activated in all four experiments. Green, pathways activated in CSNs by kaB-RAF expression as well as in cells regenerating after peripheral axotomy, or in MNs after injury and in CSNs by SCI+NPC. Purple, PTEN and mTOR pathways are activated or suppressed in regenerating CSNs and MNs only. Blue, pathways activated in CSNs but not sensory or motor neurons. Yellow, neuroinflammatory pathways activated in all axotomy paradigms but not in the non-injured kaB-Raf expressing CSNs. Tan, pathways strongly activated in CSNs expressing kaB-RAF but not in any of the lesioned and regenerating neuron types.

| Ingenuity Canonical Pathways | CSN | DRG, SCI |  | FMN, FNI |  | CSN, SCI+NPC |  |
| --- | --- | --- | --- | --- | --- | --- | --- |
|  | ckaB-RAF | 1d | 4d | 1d | 4d | 3d | 7d |
| CREB Signaling in Neurons | 2.887 | 4.841 | 2.236 | 2.236 | 2.411 | 4.667 | 4.596 |
| G-Protein Coupled Receptor Signaling | 4.003 | 4.003 |  |  | 3.213 | 3.363 | 2.556 |
| FAK Signaling | 4.622 | 4.522 | 2.111 | 2.066 | 3.286 | 5.555 | 4.802 |
| STAT3 Pathway | 2.53 | 2.53 | 2 | 2.668 |  |  |  |
| IL-6 Signaling |  | 2.121 | 3 | 2.524 | 2.121 |  |  |
| Signaling by Rho Family GTPases | 2.53 |  |  | 2.5 |  |  |  |
| Ephrin Receptor Signaling | 2.84 |  |  |  | 2.041 | 2 |  |
| PTEN Signaling |  |  |  |  |  | -2.449 |  |
| mTOR Signaling |  |  |  | 3.357 | 2.2 |  | 2.646 |
| Integrin Signaling | 2.496 |  |  |  |  | 2.84 | 2.828 |
| cAMP-mediated Signaling | 2.121 |  |  |  |  | 2.887 | 2.309 |
| FGF Signaling | 2.236 |  |  |  |  | 2.236 | 2.236 |
| Neuroinflammation Signaling Pathway |  | 2.673 | 3.207 | 2.353 | 3.677 | 2.524 | 2.309 |
| Synaptogenesis Signaling Pathway | 3.441 |  |  |  |  |  |  |
| Calcium Signaling | 2.121 |  |  |  |  |  |  |
| NGF Signaling | 2.121 |  |  |  |  |  |  |

**Video S1.** Whole mount staining and 3D imaging of adult spinal cord interneurons. AAV2 expressing transsynaptic tracer WGACreER was unilaterally injected in the motor cortex of an 8-week-old *lsl-tdTom* mouse. 14 days after AAV injection mouse is treated with tamoxifen for 5 consecutive days.

10 days after the last tamoxifen administration, mouse was perfused with 4% PFA and the spinal cord was cleared using a iDISCO+ method and imaged using a light sheet microscope subsequently.

## References

1. J. F. Diaz Quiroz, K. Echeverri, Spinal cord regeneration: where fish, frogs and salamanders lead the way, can we follow? Biochem J 451, 353–364 (2013).

2. J. P. Rasmussen, A. Sagasti, Learning to swim, again: Axon regeneration in fish. Exp Neurol 287, 318–330 (2017).

3. J. W. Fawcett, J. Verhaagen, Intrinsic Determinants of Axon Regeneration. Dev Neurobiol 78, 890–897 (2018).

4. S. G. Varadarajan, J. L. Hunyara, N. R. Hamilton, A. L. Kolodkin, A. D. Huberman, Central nervous system regeneration. Cell 185, 77–94 (2022).

5. G. Courtine, M. V. Sofroniew, Spinal cord repair: advances in biology and technology. Nat Med 25, 898–908 (2019).

6. A. Markus, J. Zhong, W. D. Snider, Raf and akt mediate distinct aspects of sensory axon growth. Neuron 35, 65–76 (2002).

7. J. Zhong et al., Raf kinase signaling functions in sensory neuron differentiation and axon growth in vivo. Nat Neurosci 10, 598–607 (2007).

8. T. N. Vizard, M. Newton, L. Howard, S. Wyatt, A. M. Davies, ERK signaling mediates CaSR-promoted axon growth. Neurosci Lett 603, 77–83 (2015).

9. X. Li et al., MEK Is a Key Regulator of Gliogenesis in the Developing Brain. Neuron 75, 1035–1050 (2012).

10. K. J. O’Donovan et al., B-RAF kinase drives developmental axon growth and promotes axon regeneration in the injured mature CNS. J Exp Med 211, 801–814 (2014).

11. A. Blesch, M. H. Tuszynski, Spinal cord injury: plasticity, regeneration and the challenge of translational drug development. Trends Neurosci 32, 41–47 (2009).

12. R. Deumens, G. C. Koopmans, E. A. Joosten, Regeneration of descending axon tracts after spinal cord injury. Prog Neurobiol 77, 57–89 (2005).

13. D. D. Pearse et al., cAMP and Schwann cells promote axonal growth and functional recovery after spinal cord injury. Nat Med 10, 610–616 (2004).

14. O. Raineteau, M. E. Schwab, Plasticity of motor systems after incomplete spinal cord injury. Nat Rev Neurosci 2, 263–273 (2001).

15. M. Belci, M. Catley, M. Husain, H. L. Frankel, N. J. Davey, Magnetic brain stimulation can improve clinical outcome in incomplete spinal cord injured patients. Spinal Cord 42, 417–419 (2004).

16. R. Nardone et al., rTMS modulates reciprocal inhibition in patients with traumatic spinal cord injury. Spinal Cord 52, 831–835 (2014).

17. J. Gomes-Osman, E. C. Field-Fote, Improvements in hand function in adults with chronic tetraplegia following a multiday 10-Hz repetitive transcranial magnetic stimulation intervention combined with repetitive task practice. J Neurol Phys Ther 39, 23–30 (2015).

18. X. Chen, et al., High-Throughput Mapping of Long-Range Neuronal Projection Using In Situ Sequencing. Cell 179, 772–786 e719 (2019).

19. J. Liu et al., Cell-specific translational profiling in acute kidney injury. J Clin Invest 124, 1242–1254 (2014).

20. L. Madisen et al., A robust and high-throughput Cre reporting and characterization system for the whole mouse brain. Nat Neurosci 13, 133–140 (2010).

21. K. Mercer et al., Expression of endogenous oncogenic V600EB-raf induces proliferation and developmental defects in mice and transformation of primary fibroblasts. Cancer Res 65, 11493–11500 (2005).

22. M. Heiman, R. Kulicke, R. J. Fenster, P. Greengard, N. Heintz, Cell type-specific mRNA purification by translating ribosome affinity purification (TRAP). Nat Protoc 9, 1282–1291 (2014).

23. A. B. Keenan et al., ChEA3: transcription factor enrichment analysis by orthogonal omics integration. Nucleic Acids Res 47, W212–W224 (2019).

24. M. V. Kuleshov et al., Enrichr: a comprehensive gene set enrichment analysis web server 2016 update. Nucleic Acids Res 44, W90–97 (2016).

25. V. Chandran et al., A Systems-Level Analysis of the Peripheral Nerve Intrinsic Axonal Growth Program. Neuron 89, 956–970 (2016).

26. J. K. Lerch, Y. R. Martinez-Ondaro, J. L. Bixby, V. P. Lemmon, cJun promotes CNS axon growth. Mol Cell Neurosci 59, 97–105 (2014).

27. M. C. Danzi et al., The effect of Jun dimerization on neurite outgrowth and motif binding. Mol Cell Neurosci 92, 114–127 (2018).

28. M. R. J. Mason et al., The Jun-dependent axon regeneration gene program: Jun promotes regeneration over plasticity. Hum Mol Genet, (2021).

29. S. P. Dhara et al., Cellular reprogramming for successful CNS axon regeneration is driven by a temporally changing cast of transcription factors. Sci Rep 9, 14198 (2019).

30. B. K. Lee et al., Fosl1 overexpression directly activates trophoblast-specific gene expression programs in embryonic stem cells. Stem Cell Res 26, 95–102 (2018).

31. K. Mendes et al., The epigenetic pioneer EGR2 initiates DNA demethylation in differentiating monocytes at both stable and transient binding sites. Nat Commun 12, 1556 (2021).

32. M. Fernandez Garcia et al., Structural Features of Transcription Factors Associating with Nucleosome Binding. Mol Cell 75, 921–932 e926 (2019).

33. A. Essafi et al., A wt1-controlled chromatin switching mechanism underpins tissue-specific wnt4 activation and repression. Developmental cell 21, 559–574 (2011).

34. T. J. Kendall et al., Embryonic mesothelial-derived hepatic lineage of quiescent and heterogenous scar-orchestrating cells defined but suppressed by WT1. Nat Commun 10, 4688 (2019).

35. S. Li et al., The transcriptional landscape of dorsal root ganglia after sciatic nerve transection. Sci Rep 5, 16888 (2015).

36. D. L. Moore et al., KLF family members regulate intrinsic axon regeneration ability. Science 326, 298–301 (2009).

37. J. H. Xu et al., Deletion of Kruppel-like factor-4 promotes axonal regeneration in mammals. Neural Regen Res 16, 166–171 (2021).

38. M. Mahar, V. Cavalli, Intrinsic mechanisms of neuronal axon regeneration. Nat Rev Neurosci 19, 323–337 (2018).

39. F. Elsaeidi, M. A. Bemben, X. F. Zhao, D. Goldman, Jak/Stat signaling stimulates zebrafish optic nerve regeneration and overcomes the inhibitory actions of Socs3 and Sfpq. J Neurosci 34, 2632–2644 (2014).

40. Y. Lu, S. Belin, Z. He, Signaling regulations of neuronal regenerative ability. Curr Opin Neurobiol 27, 135–142 (2014).

41. P. D. Smith et al., SOCS3 deletion promotes optic nerve regeneration in vivo. Neuron 64, 617–623 (2009).

42. J. Zhong, I. D. Dietzel, P. Wahle, M. Kopf, R. Heumann, Sensory impairments and delayed regeneration of sensory axons in interleukin-6-deficient mice. J Neurosci 19, 4305–4313 (1999).

43. K. K. Park et al., Promoting axon regeneration in the adult CNS by modulation of the PTEN/mTOR pathway. Science 322, 963–966 (2008).

44. K. Liu et al., PTEN deletion enhances the regenerative ability of adult corticospinal neurons. Nat Neurosci 13, 1075–1081 (2010).

45. F. Sun et al., Sustained axon regeneration induced by co-deletion of PTEN and SOCS3. Nature 480, 372–375 (2011).

46. F. M. Bareyre et al., In vivo imaging reveals a phase-specific role of STAT3 during central and peripheral nervous system axon regeneration. Proc Natl Acad Sci U S A 108, 6282–6287 (2011).

47. G. H. D. Poplawski et al., Injured adult neurons regress to an embryonic transcriptional growth state. Nature 581, 77–82 (2020).

48. M. H. Tuszynski, O. Steward, Concepts and methods for the study of axonal regeneration in the CNS. Neuron 74, 777–791 (2012).

49. V. Dietz, K. Fouad, Restoration of sensorimotor functions after spinal cord injury. Brain 137, 654–667 (2014).

50. D. Jin et al., Restoration of skilled locomotion by sprouting corticospinal axons induced by co-deletion of PTEN and SOCS3. Nat Commun 6, 8074 (2015).

51. M. L. Starkey, C. Bleul, I. C. Maier, M. E. Schwab, Rehabilitative training following unilateral pyramidotomy in adult rats improves forelimb function in a non-task-specific way. Exp Neurol 232, 81–89 (2011).

52. E. Azim, J. Jiang, B. Alstermark, T. M. Jessell, Skilled reaching relies on a V2a propriospinal internal copy circuit. Nature 508, 357–363 (2014).

53. M. L. Starkey et al., Assessing behavioural function following a pyramidotomy lesion of the corticospinal tract in adult mice. Exp Neurol 195, 524–539 (2005).

54. Z. Gu et al., Control of species-dependent cortico-motoneuronal connections underlying manual dexterity. Science 357, 400–404 (2017).

55. E. S. Rosenzweig, J. W. McDonald, Rodent models for treatment of spinal cord injury: research trends and progress toward useful repair. Curr Opin Neurol 17, 121–131 (2004).

56. B. J. Cummings, C. Engesser-Cesar, G. Cadena, A. J. Anderson, Adaptation of a ladder beam walking task to assess locomotor recovery in mice following spinal cord injury. Behav Brain Res 177, 232–241 (2007).

57. Y. Liu et al., A Sensitized IGF1 Treatment Restores Corticospinal Axon-Dependent Functions. Neuron 95, 817–833 e814 (2017).

58. M. Hirano et al., Highly efficient retrograde gene transfer into motor neurons by a lentiviral vector pseudotyped with fusion glycoprotein. PLoS One 8, e75896 (2013).

59. L. B. Rosen, D. D. Ginty, M. J. Weber, M. E. Greenberg, Membrane depolarization and calcium influx stimulate MEK and MAP kinase via activation of Ras. Neuron 12, 1207–1221 (1994).

60. S. Kupzig, S. A. Walker, P. J. Cullen, The frequencies of calcium oscillations are optimized for efficient calcium-mediated activation of Ras and the ERK/MAPK cascade. P Natl Acad Sci USA 102, 7577–7582 (2005).

61. R. Yasuda et al., Supersensitive Ras activation in dendrites and spines revealed by two-photon fluorescence lifetime imaging. Nat Neurosci 9, 283–291 (2006).

62. S. N. Makarov, G. M. Noetscher, T. Raij, A. Nummenmaa, A Quasi-Static Boundary Element Approach With Fast Multipole Acceleration for High-Resolution Bioelectromagnetic Models. IEEE Trans Biomed Eng 65, 2675–2683 (2018).

63. J. Parthoens et al., Performance Characterization of an Actively Cooled Repetitive Transcranial Magnetic Stimulation Coil for the Rat. Neuromodulation 19, 459–468 (2016).

64. B. Lafon, A. Rahman, M. Bikson, L. C. Parra, Direct Current Stimulation Alters Neuronal Input/Output Function. Brain Stimul 10, 36–45 (2017).

65. W. H. Lee, S. H. Lisanby, A. F. Laine, A. V. Peterchev, Electric Field Model of Transcranial Electric Stimulation in Nonhuman Primates: Correspondence to Individual Motor Threshold. IEEE Trans Biomed Eng 62, 2095–2105 (2015).

66. C. Marques et al., NF1 regulates mesenchymal glioblastoma plasticity and aggressiveness through the AP-1 transcription factor FOSL1. Elife 10, (2021).

67. G. Raivich et al., The AP-1 transcription factor c-Jun is required for efficient axonal regeneration. Neuron 43, 57–67 (2004).

68. C. A. Ruff et al., Neuronal c-Jun is required for successful axonal regeneration, but the effects of phosphorylation of its N-terminus are moderate. J Neurochem 121, 607–618 (2012).

69. I. Venkatesh, M. T. Simpson, D. M. Coley, M. G. Blackmore, Epigenetic profiling reveals a developmental decrease in promoter accessibility during cortical maturation in vivo. Neuroepigenetics 8, 19–26 (2016).

70. S. Qin, Y. Zou, C. L. Zhang, Cross-talk between KLF4 and STAT3 regulates axon regeneration. Nat Commun 4, 2633 (2013).

71. K. Kadoya et al., Spinal cord reconstitution with homologous neural grafts enables robust corticospinal regeneration. Nat Med 22, 479–487 (2016).

72. K. Tong et al., Degree of Tissue Differentiation Dictates Susceptibility to BRAF-Driven Colorectal Cancer. Cell Rep 21, 3833–3845 (2017).

73. Y. Chen, Y. He, M. J. DeVivo, Changing Demographics and Injury Profile of New Traumatic Spinal Cord Injuries in the United States, 1972-2014. Arch Phys Med Rehabil 97, 1610–1619 (2016).

74. C. S. Siegel, K. L. Fink, S. M. Strittmatter, W. B. Cafferty, Plasticity of intact rubral projections mediates spontaneous recovery of function after corticospinal tract injury. J Neurosci 35, 1443–1457 (2015).

75. L. Asboth et al., Cortico-reticulo-spinal circuit reorganization enables functional recovery after severe spinal cord contusion. Nat Neurosci 21, 576–588 (2018).

76. N. Zareen et al., Stimulation-dependent remodeling of the corticospinal tract requires reactivation of growth-promoting developmental signaling pathways. Exp Neurol 307, 133–144 (2018).

77. E. S. Rosenzweig et al., Extensive spontaneous plasticity of corticospinal projections after primate spinal cord injury. Nat Neurosci 13, 1505–1510 (2010).

78. F. B. Wagner et al., Targeted neurotechnology restores walking in humans with spinal cord injury. Nature 563, 65–71 (2018).

79. N. Jayaprakash et al., Optogenetic Interrogation of Functional Synapse Formation by Corticospinal Tract Axons in the Injured Spinal Cord. J Neurosci 36, 5877–5890 (2016).

80. Z. Wang, A. Reynolds, A. Kirry, C. Nienhaus, M. G. Blackmore, Overexpression of Sox11 promotes corticospinal tract regeneration after spinal injury while interfering with functional recovery. J Neurosci 35, 3139–3145 (2015).

81. M. Fujiki, K. M. Yee, O. Steward, Non-invasive High Frequency Repetitive Transcranial Magnetic Stimulation (hfrTMS) Robustly Activates Molecular Pathways Implicated in Neuronal Growth and Synaptic Plasticity in Select Populations of Neurons. Front Neurosci 14, 558 (2020).

82. K. Makowiecki, A. R. Harvey, R. M. Sherrard, J. Rodger, Low-intensity repetitive transcranial magnetic stimulation improves abnormal visual cortical circuit topography and upregulates BDNF in mice. J Neurosci 34, 10780–10792 (2014).

83. M. B. Muller, N. Toschi, A. E. Kresse, A. Post, M. E. Keck, Long-term repetitive transcranial magnetic stimulation increases the expression of brain-derived neurotrophic factor and cholecystokinin mRNA, but not neuropeptide tyrosine mRNA in specific areas of rat brain. Neuropsychopharmacology 23, 205–215 (2000).

84. M. Niimi et al., Role of Brain-Derived Neurotrophic Factor in Beneficial Effects of Repetitive Transcranial Magnetic Stimulation for Upper Limb Hemiparesis after Stroke. PLoS One 11, e0152241 (2016).

85. R. Gersner, E. Kravetz, J. Feil, G. Pell, A. Zangen, Long-term effects of repetitive transcranial magnetic stimulation on markers for neuroplasticity: differential outcomes in anesthetized and awake animals. J Neurosci 31, 7521–7526 (2011).

86. M. S. Kim et al., Efficacy of cumulative high-frequency rTMS on freezing of gait in Parkinson’s disease. Restor Neurol Neurosci 33, 521–530 (2015).

87. W. H. Chang et al., Long-term effects of rTMS on motor recovery in patients after subacute stroke. J Rehabil Med 42, 758–764 (2010).

88. N. Brihmat et al., High-Frequency rTMS Combined with Task-Specific Hand Motor Training Modulates Corticospinal Plasticity in Motor Complete Spinal Cord Injury: A case report. Conf Proc IEEE Eng Med Biol Soc **Accpeted**, (2022).

89. K. Howe et al., The zebrafish reference genome sequence and its relationship to the human genome. Nature 496, 498–503 (2013).

90. V. Bissonauth, S. Roy, M. Gravel, S. Guillemette, J. Charron, Requirement for Map2k1 (Mek1) in extra-embryonic ectoderm during placentogenesis. Development 133, 3429–3440 (2006).

91. L.-F. Bélanger et al., Mek2 Is dispensable for mouse growth and development. Mol. Cell. Biol. 23, 4778–4787 (2003).

92. R. Lesche et al., Cre/loxP-mediated inactivation of the murine Pten tumor suppressor gene. Genesis 32, 148–149 (2002).

93. F. Boato et al., C3 peptide enhances recovery from spinal cord injury by improved regenerative growth of descending fiber tracts. J Cell Sci 123, 1652–1662 (2010).

94. C. A. Danilov, O. Steward, Conditional genetic deletion of PTEN after a spinal cord injury enhances regenerative growth of CST axons and motor function recovery in mice. Exp Neurol 266, 147–160 (2015).

95. K. Zukor et al., Short hairpin RNA against PTEN enhances regenerative growth of corticospinal tract axons after spinal cord injury. J Neurosci 33, 15350–15361 (2013).

96. X. Wang et al., Deconstruction of Corticospinal Circuits for Goal-Directed Motor Skills. Cell 171, 440–455 e414 (2017).

97. N. Renier et al., iDISCO: a simple, rapid method to immunolabel large tissue samples for volume imaging. Cell 159, 896–910 (2014).

98. H. L. Feng, L. Yan, Y. Z. Guan, L. Y. Cui, Effects of transcranial magnetic stimulation on motor cortical excitability and neurofunction after cerebral ischemia-reperfusion injury in rats. Chin Med Sci J 20, 226–230 (2005).

99. M. Sykes et al., Differences in Motor Evoked Potentials Induced in Rats by Transcranial Magnetic Stimulation under Two Separate Anesthetics: Implications for Plasticity Studies. Front Neural Circuits 10, 80 (2016).

100. B. Dogdas, D. Stout, A. F. Chatziioannou, R. M. Leahy, Digimouse: a 3D whole body mouse atlas from CT and cryosection data. Phys Med Biol 52, 577–587 (2007).

101. A. Nummenmaa et al., Comparison of spherical and realistically shaped boundary element head models for transcranial magnetic stimulation navigation. Clin Neurophysiol 124, 1995–2007 (2013).

102. P. C. Miranda, M. Hallett, P. J. Basser, The electric field induced in the brain by magnetic stimulation: a 3-D finite-element analysis of the effect of tissue heterogeneity and anisotropy. IEEE Trans Biomed Eng 50, 1074–1085 (2003).

103. R. Willenberg, K. Zukor, K. Liu, Z. He, O. Steward, Variable laterality of corticospinal tract axons that regenerate after spinal cord injury as a result of PTEN deletion or knock-down. J Comp Neurol 524, 2654–2676 (2016).

104. A. Kramer, J. Green, J. Pollard, Jr., S. Tugendreich, Causal analysis approaches in Ingenuity Pathway Analysis. Bioinformatics 30, 523–530 (2014).

